# Multidisciplinary assessments of a potentially novel acute tissue loss disease on Florida *Orbicella* corals

**DOI:** 10.64898/2025.12.10.693521

**Authors:** Karen L. Neely, Allison Cauvin, Yasunari Kiryu, Erin Papke, Grace E. Kennedy, Arelys A. Chaparro, Kalie Januskiewicz, Esther C. Peters, Lindsay K. Huebner, Jan H. Landsberg, Blake Ushijima, Julie L. Meyer

## Abstract

Coral disease outbreaks are an increasingly common threat to reefs. While coral disease research is expanding rapidly, there are still monumental challenges in diagnosing and differentiating among the different diseases on a reef. We used a collaborative multidisciplinary approach to characterize a potentially novel disease affecting *Orbicella faveolata* colonies in the Florida Keys. We tagged and fate-tracked individual lesions for progression and cessation rates, reanalyzed in situ photographs of previously monitored corals to determine prevalence and seasonality, and assessed the efficacy of amoxicillin treatments. Samples collected from lesions as well as unaffected and healthy controls were used to examine microbiomes, assess nine coral pathological parameters via histology, and measure 19 symbiont-specific physiological metrics using transmission electron microscopy (TEM). Across all analyses, we concluded that the observed disease is unlikely to be SCTLD. Many, but not all, metrics had similarities to previous descriptions of white plague, and so we additionally conducted preliminary histology on presumed white plague samples of *O. franksi* for comparison. However, the dearth of quantifiable histology and TEM studies on white plague did not allow us to conclusively confirm or refute comparisons to white plague. Using the recommended coral disease nomenclature, we define this specific outbreak as “*Orbicella* acute tissue loss disease” (OATLD). We provide unprecedented, quantified descriptions across numerous metrics of both diseased and control colonies. We suggest that these data lay the groundwork for future efforts on this disease as well as a comprehensive set of parameters against which other diseases can be compared.

## Introduction

Scleractinian (stony) corals are susceptible to a variety of stressors, but one of the most common and potentially devastating is disease. Disease outbreaks can cause acute and mass mortalities of corals. The number of reported coral diseases has been steadily increasing (Richardson, 1998; Harvell et al., 1999), and the incidence of disease outbreaks is predicted to increase over time (Maynard et al., 2015; Burke et al., 2023). The Caribbean basin is a hot spot for coral diseases, and major region-wide coral diseases such as white band disease (Antonius, 1981; Gladfelter, 1982) and stony coral tissue loss disease (SCTLD) (Precht et al., 2016; Hawthorn et al., 2024) have resulted in the widespread loss of corals, including near regional extinction of some species (Gladfelter, 1982; Neely et al., 2021a).

Differentiating among coral diseases in situ presents various challenges. The number of disease signs that corals can present is limited, and the appearance of disease lesions can exhibit variations over time as well as both intra- and inter-specifically (Florida Coral Disease Response Research & Epidemiology Team, 2018; Aeby et al., 2021). Additionally, the etiological agents for most coral diseases have not been identified (reviewed in Vega Thurber et al. (2020) and Richardson (1998)), severely hindering diagnostic tool development. Nevertheless, the identification of etiological agents responsible for coral diseases remains important for monitoring populations, tracking outbreaks, mitigating transmission sources, and developing disease treatments. To better aid field identification, the Coral Disease and Health Consortium has proposed standardized nomenclature for lesion appearance and distribution across colonies (Rogers, 2010). Additionally, laboratory tools to better identify coral diseases continue to be improved. These include microbiome analyses, histology, and transmission electron microscopy (TEM) imaging (Pollock et al., 2011; Hawthorn et al., 2023; Papke et al., 2024a; Papke et al., 2024b).

Here, we describe a collaborative multidisciplinary approach to characterize a potentially novel disease affecting a prominent coral species in the Florida Keys, *Orbicella faveolata.* Initial investigations of this coral disease colloquially called these lesions “fast lesion progression (FLP)” (Neely, 2023b), but following the terminology laid out by the Coral Disease and Health Consortium, we now present a more accurate nomenclature: “*Orbicella* acute tissue loss disease” (OATLD). We assessed this potentially novel disease through a collaborative consortium by reexamining photos from fate-tracked *O. faveolata* colonies since 2019 for prevalence and seasonality, tracking individual lesions for progression rates and halting, assessing the efficacy of amoxicillin treatments on lesions, and testing lesions for the presence of *Vibrio coralliilyticus*. We also analyzed samples taken from OATLD-affected and healthy control corals to assess microbiomes, examine pathological parameters through histology, and assess algal endosymbiont metrics through TEM.

## Methods

### Field Assessments

#### Lesion tracking

We regularly monitored OATLD-affected corals at three sites within the Florida Keys National Marine Sanctuary: Looe Key, the Grecian/KLDR (Key Largo Dry Rocks) paired reefs, and the paired Carysfort (Carysfort South and Carysfort Main) reefs (Figure 1). Corals with presumed OATLD were tagged and mapped, measured for straight line length, width, and height, assessed for percent live, recently dead (i.e., bright white uncolonized skeleton, corallite structure intact), and old dead coverage (i.e., non-white skeleton with degraded corallite structure), and photographed. On active lesions, two nails were placed on the live/dead tissue margin 10 cm apart from each other (Neely, 2024). For lesions longer than 50 cm, multiple nail sets were placed along the lesion line, each at least 20 cm apart from the neighboring set.

**Fig. 1:**
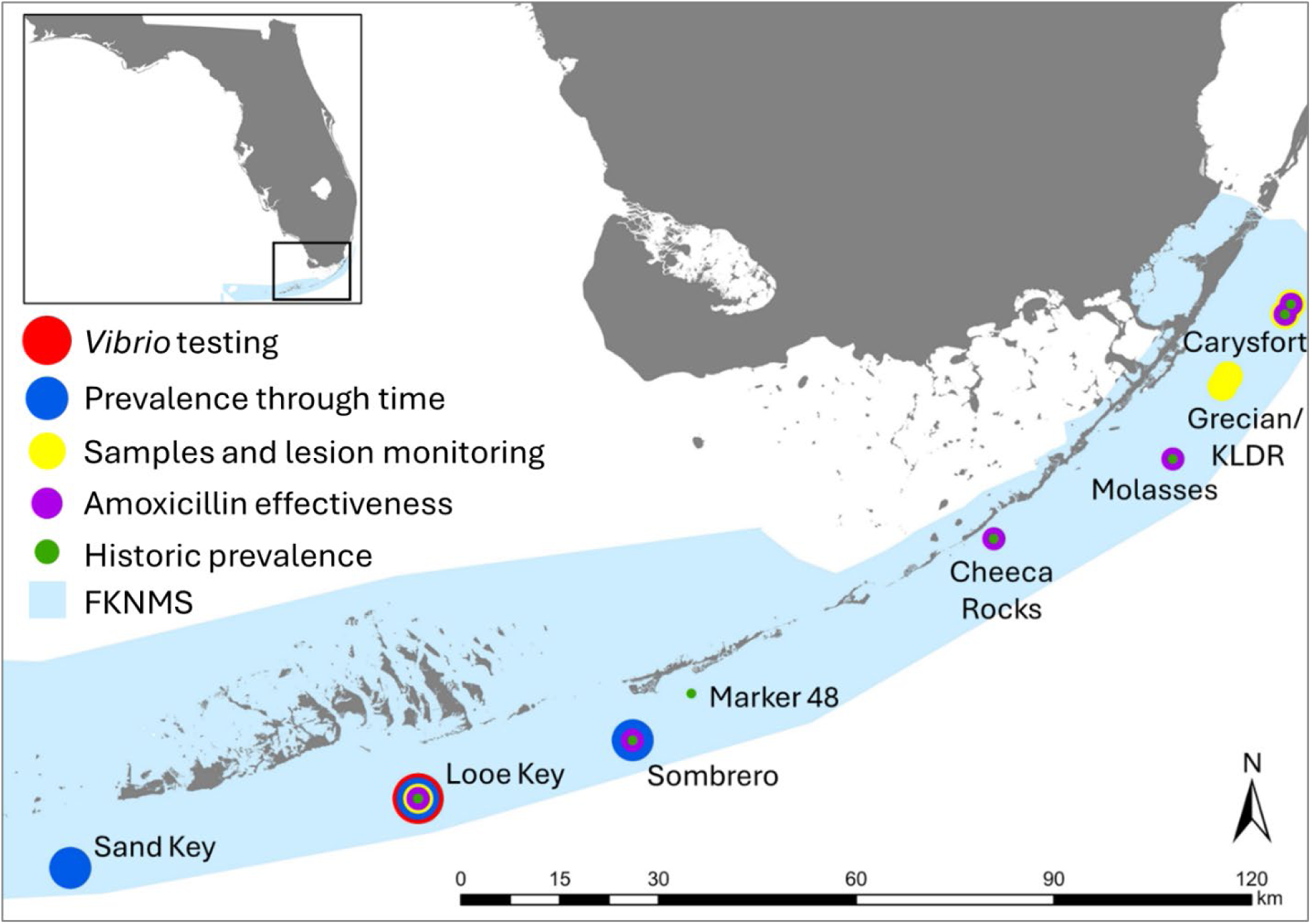
Map of reef sites used for various assessments of OATLD-affected *Orbicella faveolata* colonies. All sites are within the Florida Keys National Marine Sanctuary (FKNMS).

For lesion tracking, each site was visited approximately every two months from January 2024 to January 2025. During each visit, any newly affected corals with measurable lesions were tagged and measured as described above. Any previously tagged corals were assessed for percent live, old dead, and recently dead. Any previous lesions were assessed as active (defined as bright white skeleton adjacent to lesion), halted, or no longer present because all previously live tissue ahead of the lesion was dead. Lesion progression rates for active lesions were measured as the distance the lesion had progressed perpendicular to the marker nails divided by the number of days since the previous observation (cm / day). Measurements were taken with a flexible measuring tape laid along the coral surface to account for rugosity.

We obtained a total of 122 OATLD lesion measurements from 60 affected corals across the three sites. This included 29 colonies at Looe Key, 17 at Grecian/KLDR, and 14 at Carysfort. Proportions of halted lesions between sites were compared using a χ^2^ test. Lesion progression rates between sites were compared using a one-way ANOVA.

#### Historical presence

We used archived underwater field photographs from 2019 to 2023 to determine whether OATLD was present at Florida Keys reef sites in past years. Photographs were taken as a component of the intervention response to the SCTLD outbreak. As part of that response, over 5,000 corals in the Florida Keys National Marine Sanctuary were treated for disease (Neely et al., 2021b; Neely et al., 2025). Treatments included a topical amoxicillin paste, chlorinated epoxy, topical antimicrobials, and/or probiotics. During the SCTLD outbreak, it was presumed that any similarly-appearing disease lesions were SCTLD and treated accordingly. For each treated coral, photos were taken before the initial treatments, and these historical images were reassessed to determine to what extent lesions now presumed to be OATLD were present during past years.

At each of six reef sites (offshore: Looe Key, Sombrero, Molasses, and Carysfort. Inshore: Marker 48 and Cheeca Rocks. Figure 1), the initial photos from the first 40 *O. faveolata* colonies treated for SCTLD were reexamined to determine if corals in fact had OATLD-style lesions. OATLD lesions were defined as linear, smooth, and acute; SCTLD lesions were defined as irregular and with an undulating border. Because not all sites were first treated at the same time, and because *O. faveolata* were more common at some sites than others, the timing for the first 40 colonies treated was not consistent across sites. For example, the Lower/Middle Keys sites of Looe Key and Sombrero were first treated in 2019 when SCTLD was detected relatively early, thus all 40 of the assessments at those sites are from 2019. In contrast, the Upper Keys sites Carysfort and Molasses had already suffered substantial losses to SCTLD when treatments began, and so first appearances of disease on *O. faveolata* colonies range across multiple years. By expanding the range of years available for assessment, we were able to assess 40 *O. faveolata* colonies each from all sites except Molasses, at which only 34 *O. faveolata* colonies qualified for assessment. We compared the proportions of corals exhibiting OATLD lesions between sites with a χ ^2^ test followed by pairwise χ ^2^ tests between sites.

#### Amoxicillin effectiveness

A topical amoxicillin paste (98% amoxicillin trihydrate (Phytotech) mixed with Base2b (Ocean Alchemists) in a 1:8 by weight ratio) was used to treat SCTLD lesions across numerous reef sites in the Florida Keys, with lesions and corals fate-tracked approximately every two months for up to five years after application (Neely et al., 2021b; Neely et al., 2025). Using the historical photographs outlined above from corals that were presumed to have OATLD lesions rather than SCLTD lesions, we assessed the efficacy of the amoxicillin treatment. From corals at five reef sites (Looe Key, Sombrero, Cheeca Rocks, Molasses, and Carysfort) we assessed 40 suspected OATLD lesions treated in 2019 to 2020, and 23 lesions treated in 2022 to 2023. Treatment efficacy was assessed by comparing the initial treatment photos with lesion photos from the subsequent monitoring (no more than three months after treatment). Effective treatment was defined as the lesion halting at the treatment line, while ineffective treatment was defined as the lesion continuing past the treatment line. The proportion of OATLD lesions that responded to treatments were compared using generalized linear models assessing site, time period (2019-2020 vs 2022-2023), and the interaction between the two. We chose the best model based on Akaike Information Criterion scores (Akaike, 1973) and conducted emmeans post-hoc tests (Lenth, 2018).

#### Seasonal assessments

At three offshore sites with large numbers of *O. faveolata* colonies (Sand Key = 130, Looe Key = 715, and Sombrero = 78; Figure 1), photographs of each *O. faveolata* colony from each monitoring period (2019/20 – 2023) were assessed to identify whether any lesions presented as SCTLD, OATLD, or both. Because each site was fully surveyed (divers swam the entire reef area) during every monitoring period, and all newly diseased corals were tagged and subsequently monitored, we made the assumption that the number of *O. faveolata* colonies at the conclusion of monitoring for SCTLD (July 2024) was the total number of known susceptible colonies to SCLTD and/or OATLD. For each monitoring period, we divided the number of colonies affected by OATLD by this total number of colonies. However, if a colony died or went missing (generally following hurricanes), we excluded it from the denominator (total number) for the month of death and all subsequent months. For each monitoring period, we also excluded any colonies that were specifically noted as not found during that monitoring event. If a colony was affected with OATLD during multiple monitoring periods, it was included in the numerator for all affected periods.

To determine if there was a seasonality pattern to the prevalence of OATLD on reefs, the months that colonies were affected with OATLD were first converted to angles to prepare the data for circular uniformity tests (Landler et al., 2018). Rayleigh tests of uniformity were then performed on the data to determine if there was a significant seasonality component. Analyses were conducted using the *circular* package (Lund et al., 2017) in R (R Core Team, 2024).

#### Colony Size

For colonies at Sand Key, Looe Key, and Sombrero, we also assessed whether colony size affected the probability of developing OATLD. We compared the maximum dimension of *O. faveolata* colonies affected with presumed OATLD at least once in their monitoring history to the maximum dimension of other tagged *O. faveolata* colonies never recorded to be affected by OATLD. Because maximum diameters were not normally distributed (Shapiro test), we used Wilcoxon-Rank tests (Wilcoxon, 1945) to compare the two groups.

#### Detection of the VcpA metalloprotease

At Looe Key, we tested five apparent OATLD lesions and five apparent SCTLD lesions in December 2022 for the presence of the VcpA metalloprotease produced by *Vibrio coralliilyticus*. This bacterium can be associated with coral diseases and is known to exacerbate the progression rates of SCTLD (Ushijima et al., 2020). We used 10 cc needle-less syringes to agitate the lesion margin and collect the mucus/tissue slurry, which were then screened with the VcpA RapidTests described in Ushijima et al. (2020). We compared the proportion of positive results between the OATLD and SCTLD lesions using a Fisher’s Exact test.

### Sampling protocol

Samples for microbiome analyses, histology, and TEM were taken from *O. faveolata* colonies at three sites: Looe Key, Grecian/KLDR, and Carysfort (Figure 1). All samples were taken between January 28 to 31, 2024. At each reef site, five colonies with active OATLD lesions and three apparently healthy colonies were selected (Supplemental Table 1). A full sample set consisted of samples for microbiome analyses, histology, and TEM, all taken directly adjacent to each other. From each apparently healthy colony, one full set of samples was collected (healthy: H). From each diseased colony, two full sample sets were collected: one from affected tissue directly adjacent to but not encompassing the live/dead tissue margin on the disease lesion (diseased; D) and one from apparently unaffected tissue at least 30 cm from any active OATLD lesions (unaffected; U). Additional samples were taken for microbiome analyses. Seven additional apparently healthy and three additional OATLD-affected colonies were selected, for a total of 10 healthy and 10 diseased corals at Looe Key and Grecian/KLDR. OATLD-affected colonies were not abundant at Carysfort during the sampling time period, and so a total of 10 apparently healthy and 5 diseased corals were sampled for microbiome at that site.

#### Microbiome

A total of 105 samples were collected from 55 colonies for microbiome analyses. At each of the three sites, a single sample was taken from each of 10 H colonies. At Looe Key and Grecian/KLDR, samples were also taken from each of 10 OATLD-affected colonies, with two syringes from each colony’s D area and one syringe from each colony’s U area. At Carysfort, only 5 diseased colonies could be found, with D and U samples taken accordingly. Mucus-tissue slurry samples were collected by scraping the coral tissue with a sterile syringe tip and using the syringe to draw up released mucus and tissue. All samples were sent on ice overnight to the University of Florida (UF) for processing and 16S rRNA gene sequencing.

#### Histology

A total of 39 *O. faveolata* tissue/skeleton biopsy core samples were collected from 24 colonies for histology. At each of the three sites, 5 D samples, 5 U samples, and 3 H samples were collected using 2.5 cm diameter hole punches. Samples were fixed with 20% Z-Fix (Anatech Ltd.) seawater solution and transported to the Florida Fish and Wildlife Conservation Commission – Fish and Wildlife Research Institute (FWC-FWRI) Histology Laboratory, St. Petersburg, FL.

#### TEM

A total of 63 TEM samples were collected from 24 colonies. At each of three sites, 2 cores from 5 D areas, 1 core from 5 U areas, and 2 cores from 3 H were extracted from the colonies using a 1.5 cm diameter hole punch. Samples were fixed and stored according to Papke et al. (2024b). The fixative contained 2.5% glutaraldehyde and 2% paraformaldehyde prepared in Instant Ocean (pH 8, 35 ppt), and samples were stored at 4°C and shipped to the University of North Carolina Wilmington (UNCW).

### Microbiome Assessments

#### Molecular Methods

A total of 77 samples were assessed: Carysfort = 10 H, 5 U, 5 D; Grecian/KLDR = 8 H, 10 U, 10 D; Looe Key= 10 H, 10 U, 9 D. For each of the coral mucus-tissue slurries, DNA was extracted using a Qiagen DNeasy PowerBiofilm Kit followed by a Qiagen DNeasy PowerClean Pro Cleanup Kit. Two reagent blanks were generated and processed identically to the biological samples. The V4 hypervariable region of the 16S rRNA gene was amplified using the Earth Microbiome protocol (Caporaso et al., 2012) and primers 515F (Parada et al., 2016) and 806R (Apprill et al., 2015) as previously described in Meyer et al. (2019). Pooled amplicons were submitted for sequencing on an Illumina MiSeq with 2 x 150 bp reads (v2 cycle format) at the University of Florida Interdisciplinary Center for Biotechnology Research NextGen Sequencing core (RRID:SCR 019152).

#### Bioinformatics Methods

Adapter and primer sequences were removed from raw sequencing reads using cutadapt v.3.4 (Martin, 2011), and subsequent analyses were conducted in R (R Core Team, 2024). Read trimming, chimera removal, error correction, and ASV calling were conducted with DADA2 v.1.32.0 (Callahan et al., 2016). Taxonomy was determined with the SILVA v138.1 database (Yilmaz et al., 2013). Mitochondrial and chloroplast sequences were removed with phyloseq v.1.48.0 (McMurdie and Holmes, 2013). The decontam package v.1.24.0 (Davis et al., 2018) was used to remove potential contaminant sequences identified in reagent blanks with the prevalence-based method of contaminant identification and a threshold of 0.5. The dataset was further filtered to remove ASVs detected in less than 2 samples and at less than 0.1% relative abundance. Reads were center-log ratio transformed using the zcompositions package v.1.5.0-5 (Palarea-Albaladejo and Martín-Fernández, 2015) and Aitchison distance was calculated using the CoDaSeq package v.0.99.7 (Gloor et al., 2016) for principal components analysis.

Variance of microbial community structure was determined via permutational analysis of variance (PERMANOVA) using the adonis2 function of the vegan package v.2.7-1 (Oksanen et al., 2001). Beta diversity dispersion was calculated as distance to centroid for each health condition with the betadisper function of vegan and statistically compared by Kruskal-Wallis H tests. Dunn’s post-hoc tests with Bonferroni corrections were used when p < 0.05 to determine significant pairwise comparisons. Alpha diversity metrics, including richness, Chao1, and Shannon diversity were calculated in phyloseq and statistically compared by Kruskal-Wallis H tests after a Shapiro test for normality. The clamtest function in vegan was used to perform the Classification Method for Habitat Specialists and Generalists test (Chazdon et al., 2011) to determine which ASVs were specialized to disease lesions, specialized to healthy colonies, were generalists, or were too rare to classify using a supermajority threshold of 2/3. DESeq2 v.1.44.0 (Love et al., 2014) with the parametric Wald test (Wald, 1943) was used to determine which taxa were differentially abundant in disease lesions compared to healthy coral tissue from healthy colonies and healthy tissue from diseased colonies. P-values were adjusted using the Benjamini-Hochberg procedure (Benjamini and Hochberg, 1995). An indicator species analysis with 999 permutations was performed with the indicspecies package v.1.8.0 (Cáceres and Legendre, 2009) to identify bioindicators for OATLD disease lesions. The BLASTN algorithm (Camacho et al., 2009) was used to match OATLD-associated ASVs recovered from this study to a custom database with previously identified SCTLD-associated ASVs (Meyer et al., 2019; Becker et al., 2022). All code for data analysis and figure generation can be found at https://github.com/meyermicrobiolab/OATLD_Data_Analysis_2. Raw sequencing files are publicly available in NCBI under BioProject PRJNA1120359.

### Histology Assessments

All of the samples collected were processed and analyzed (Supplemental Table 1). Prior to histological processing, the external surface area of each sample was examined for lesions under a dissecting microscope (Leica M125 with f = 200 mm lens for low magnifications and 1.0x PLANAPO objective for high magnifications), including the presence or absence of mesenterial filaments (MFs) protruding from the surface tissue. Photomicrographs were taken using a digital camera (Jenoptik Gryphax) attached to the dissecting scope for all samples at low and high magnifications. Subsequently, all samples were decalcified in 10% ethylenediaminetetraacetic acid (EDTA) solution (Fisher BioReagents), requiring an average of 28 days (± 3.7 SD; range 19 to 37), followed by re-examination of the decalcified tissue with a dissecting microscope and photomicrography.

Decalcified tissues were then cut into smaller portions and oriented for sectioning at both radial (cross, parallel to the polyp mouth) and sagittal (longitudinal, perpendicular to the polyp mouth) angles. Routine paraffin-embedded histologic sections were sectioned at 4 µm, stained using Mayer’s hematoxylin and eosin (H&E), thionin, and Giemsa procedures (Luna, 1968; Price and Peters, 2018), and coverslipped. Tissues were also embedded with glycol methacrylate plastic resin (JB-4; Electron Microscopy Sciences) at arbitrary angles, sectioned at 4 µm, stained using Weigert’s H&E, thionin, and periodic acid–Schiff–metanil yellow procedures (PAS-MY; Quintero-Hunter et al. (1991)), and coverslipped. Sections were examined by light microscopy (Olympus BX51 with 2x–100x PLAN S APO or PLAN FL objectives) equipped with a digital camera (Olympus DP71). Histopathological parameters were recorded for presence or absence and described separately for host anatomical tissue locations and for endosymbionts.

Fisher’s exact tests with Bonferroni-adjusted p-values were used to compare the prevalence of the gross pathological and histopathological variables, with pairwise comparisons conducted for disease conditions and sampling sites (R Core Team, 2022).

### TEM Assessments

#### TEM sample preparation and imaging

Ten samples from six coral colonies were assessed for TEM. The sample colonies were: 1 healthy coral from Carysfort (H), 1 healthy coral from Grecian/KLDR (H), 3 diseased corals from Carysfort (D and U), and 1 diseased coral from Grecian/KLDR (D and U). Samples were prepared and imaged following methods described in Papke et al. (2024b). Briefly, the fixed samples were decalcified in 10% neutral EDTA (VWR). Decalcified samples were then post-fixed with 1% osmium tetroxide (Electron Microscopy Sciences), dehydrated through an ethanol series and embedded in Spurr’s resin (Electron Microscopy Sciences). Coral tissue sections (90 nm thick) placed on formvar coated copper grids were stained with UranyLess and lead citrate before imaging (Electron Microscopy Sciences). To eliminate health state-related bias, a numerical identifier was given to samples prior to imaging. All samples were imaged at UNCW’s Richard M. Dillaman Bioimaging Facility using a FEI Tecnai Spirit Bio Twin TEM at an acceleration voltage of 80kv with a tungsten filament. Alignments were performed before images were taken with TIA software (FEI Company Version 4.7 SPI build 1455) and a CCD camera (FEI Eagle 2K).

#### ImageJ Analysis

ImageJ software (version Java 1.8.0_345) was used to analyze symbiont images (Schneider et al., 2012). Quantitative parameters were scored for each symbiont based on previously measured observations (Lohr et al., 2007; Camaya et al., 2016; Landsberg et al., 2020; Work et al., 2021; Howe-Kerr et al., 2023). In total, we analyzed 22 symbionts from the two healthy samples (H), 88 symbionts from four disease unaffected samples (U), and 64 symbionts from four diseased samples (D) (Supplemental Table 1). We assessed the following parameters for each sample:

- Density and size of symbionts: Density was calculated as the number of symbionts divided by the total gastrodermis area measured in the micrograph. Size was calculated as the area of the symbiont. A scale was set prior to measuring both the gastrodermis area and symbiont, and each area was calculated by tracing the perimeter and using the ‘Measure’ function in ImageJ.
- Proportion of symbionts exhibiting degradation: Symbiont degradation was defined as less than two organelles positively identified within the cell.
- Presence and size of any cavity within the symbionts: Cavities were defined as empty space with no organelles or cellular debris. For symbionts with a cavity, the area was calculated in ImageJ, and for those without a cavity, cavity size for the sample was defined as 0.
- Presence and size of accumulation bodies within the symbionts: An accumulation body was defined as cellular debris within a membrane. For symbionts with an accumulation body, the area was calculated in ImageJ, and for those without, the accumulation body size was defined as 0.
- Presence of degraded chloroplasts: Degraded chloroplasts were defined as those without fully intact membranes, with degraded thylakoids, with vesicles separating the thylakoids, or with swollen and/or distorted shape.
- Presence of degraded thylakoids: Thylakoid degradation was defined as indistinct or non-visible membranes.
- Presence of nuclei degradation within the symbionts: Nucleus degredation was defined as the nuclear membrane not fully intact, or the chromatin not completely condensed.
- Membrane separation from the symbiosome: Separation from the symbiosome was defined as at least one membrane surrounding the cell but significant space between the inner- and outer-most membranes.
- Number and size of lipid droplets within the symbionts: Lipid droplets were defined as circular and darkly stained structures. The perimeter of each lipid droplet was measured, and total area was calculated in ImageJ. If there were multiple lipid droplets within a symbiont, the total measured area was combined. For symbionts without lipid droplets, size was defined as 0.
- Number and size of starch granules within the symbionts: The perimeter of each starch granule was measured, and the area was calculated in ImageJ. If there were multiple starch granules within a symbiont, the total measured area was combined. For symbionts without starch granules, size was defined as 0.
- Presence of a pyrenoid, and presence of a starch sheath surrounding the pyrenoid.
- Presence of viral-like particles (VLPs): VLPs were defined as described by Work et al. (2021) and Howe-Kerr et al. (2023).
- Presence of electron dense bodies: Electron dense bodies were defined as small, darkly stained, and generally circular structures.

For each metric, proportions and sizes were compared across the symbionts within each health state. For example, the two healthy coral samples had 22 symbionts assessed between them (19 from one sample and 3 from the other). Of these, 2 had cavities, and so cavity presence of H colonies was 9% (2/22). The average cavity size for the H group is the mean of all 22 symbionts, including a “0” value for any without a cavity.

Data analyses were performed using R (R Core Team, 2024) via R Studio (R Studio Team, 2024) using R Markdown (Allaire et al., 2023). A pairwise Fisher’s exact test using the package ‘stats’ (R Core Team, 2024) was performed for categorical (presence) parameters with the Holm p-adjustment. For numerical parameters, a Shapiro-Wilks normality test was conducted alongside a Levene variance test. Ultimately, we performed Kruskal-Wallis tests using the package ‘stats’. Because of the small sample sizes and to assess trends, a Dunn test (Dinno, 2017) was performed for each pairwise comparison between health states regardless of the Kruskal-Wallis result.

#### Comparison to SCTLD study

We also compared physiological metrics from this OATLD dataset with those of symbionts from SCTLD-affected *O. faveolata* published in (Work et al., 2021). The SCTLD dataset was comprised of 45 symbionts from disease-affected (D) corals and 60 symbionts from healthy (H) corals across two coral samples per health state.

We compared seven parameters: the proportion of endosymbionts within each sample that 1) contained a cavity, 2) exhibited thylakoid degredation, 3) displayed chloroplast gigantism (defined as chloroplasts occupying more than 50% of the total cell area), 4) contained a pyrenoid, 5) contained starch granules, 6) contained stellate viral-like particles (VLPs), and 7) contained electron dense bodies.

Proportion calculations were done differently for this comparison to match those of the Work et al. (2021) SCTLD dataset. The SCTLD dataset provided only averages for each health category. Therefore, for the OATLD dataset, we converted the raw data for each sample into averages for each OATLD health group. For example, within the OATLD H sample containing three symbionts, 33.3% contained a cavity, and within the H sample with 19 symbionts, 15.8% contained a cavity, creating a “H” average of 24.6% ± 8.8 SE against which the other SCTLD and OATLD health states were compared. Kruskal-Wallis tests, followed by Dunn post-hoc analyses, were used to compare metrics among OATLD and SCTLD health states.

## Results

### Field Assessments

#### General Appearance

OATLD is characterized by large, acutely progressing lesions that appear only on *Orbicella faveolata* colonies, including some of the largest individuals on the reefs. Lesions are focal or multifocal and are typically linear, with distinct lesion edges and smooth margins (Figure 2, Supplemental Table 1). They are sometimes accompanied by an adjacent margin of loose degraded tissue along the lesion edge. Lesions are observed progressing in any direction across the colony, including horizontally, vertically, and diagonally.

**Fig. 2:**
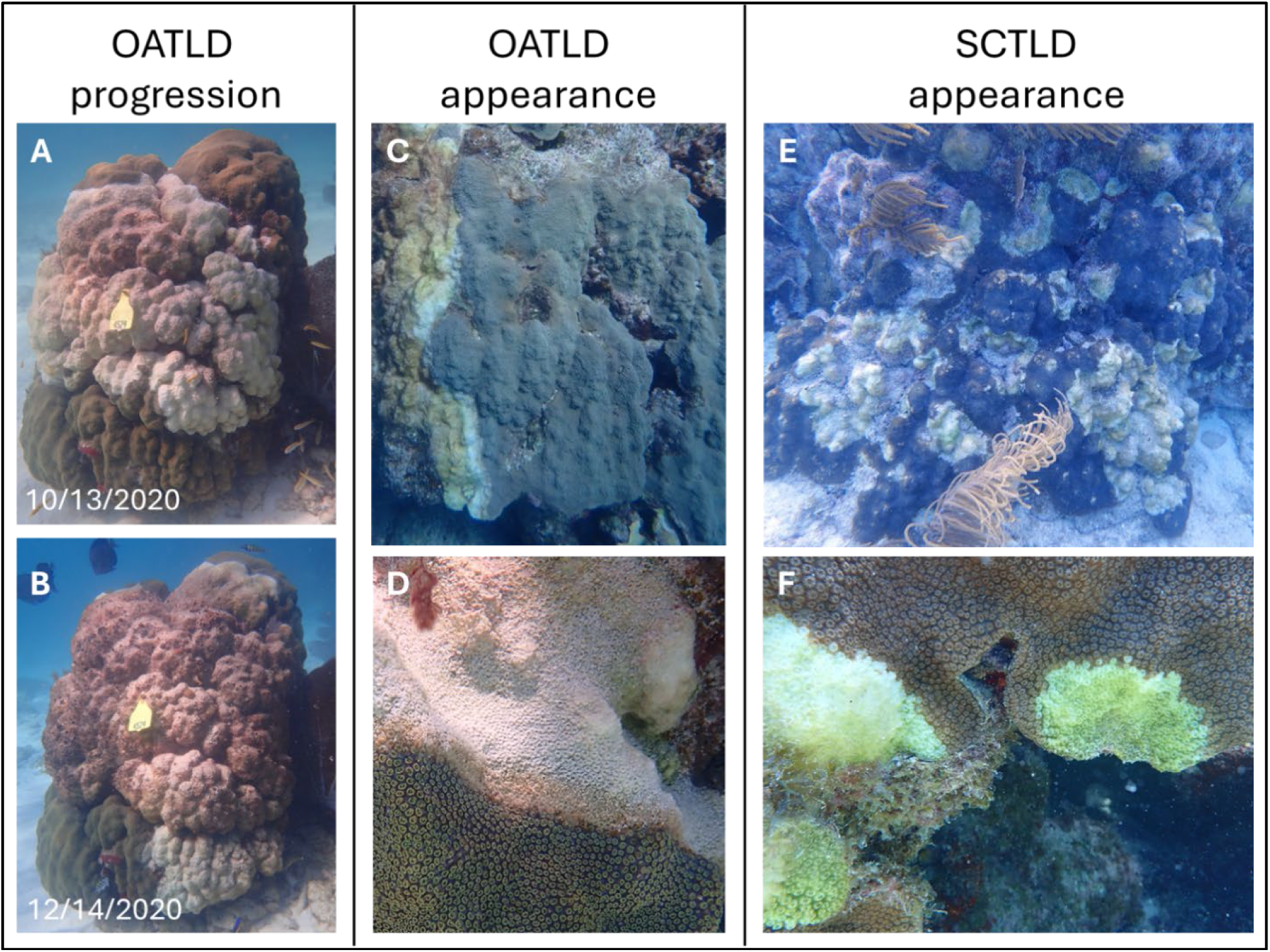
Underwater in situ photographs showing grossly visible tissue loss lesions. **(A,B)** Two-month progression of *Orbicella* acute tissue loss disease (OATLD) across an *Orbicella faveolata* colony. **(C)** Appearance of OATLD lesions showing acute linear progression and **(D)** smooth edges with an adjacent margin of degraded tissue. For comparison, **(E)** the classic appearance of stony coral tissue loss disease (SCTLD) on *O. faveolata* is multifocal, irregular, and coalescing with **(F)** an undulating border.

#### Lesion tracking

Of the 122 lesion measurements taken across all three sites, 54 (44%) halted between semi-monthly visits. There was no significant difference in the proportion of lesions halted among sites (χ^2^ ^(2, N= 122)^ = 4.8, p = 0.09. Figure 3A). Of these halted lesions, 9.3% (5/54) had 0 centimeters of lesion growth between monitoring visits, indicating that lesions halted directly after the monitoring visit. The proportion of lesions that halted did not exhibit any seasonal pattern (Figure 3B). Lesion progression rates averaged 0.43 cm/day, with a range of 0.1-1 cm per day. Progression rates were not significantly different between sites (one-way ANOVA; F = 0.37, p = 0.69. Figure 3C)

**Fig. 3:**
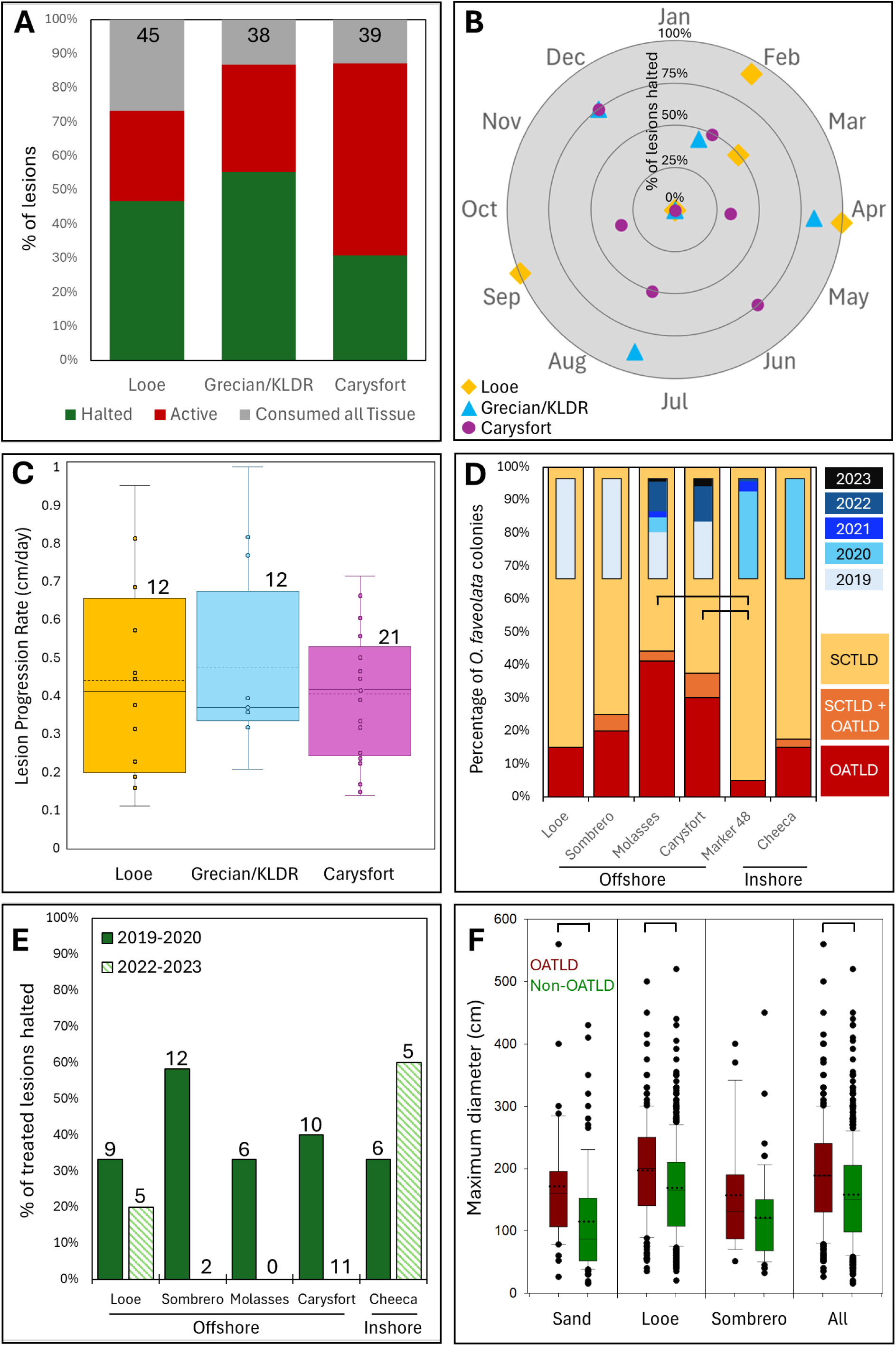
In situ observations of OATLD lesions. **(A)** Status of lesions two months after initial observation. The number of lesions assessed at each site is at the top of each column. **(B)** The percentage of previously active lesions halted during each monitoring period across the three reef sites. **(C)** Lesion progression rates. Boxes indicate the first through third quartiles, and whiskers the maximum to minimum. The median is indicated by the solid line within each box, and the mean is indicated by the dashed line. The number of lesions assessed at each site is above each box. **(D)** The proportion of the first 40 diseased colonies treated at each site (exception: Molasses N = 36) with lesions appearing as *Orbicella* acute tissue loss disease (OATLD), stony coral tissue loss disease (SCTLD), or both. The years the colonies were first observed with lesions are indicated in the smaller bars. Brackets indicate significant differences in OATLD prevalence among sites. **(E)** The percentage of OATLD lesions that halted after amoxicillin treatment across reef sites and time periods. Brackets (D,E) show significant differences. **(F)** The maximum diameter of *O. faveolata* colonies affected by OATLD (red) and those affected by SCTLD (green). Solid lines within boxes represent medians, dotted lines represent means.

#### Historical presence

OATLD was present on the first 40 *O. faveolata* (34 at Molasses) visited and treated (2019-2023) at all six monitoring sites (Figure 3D). At four of the six sites, a few corals exhibited both OATLD and SCTLD-like lesions. The proportion of colonies with OATLD lesions ranged from 5% (Marker 48) to 42% (Molasses). The proportion of OATLD-affected colonies differed significantly among sites, (χ ^2^ ^(5, N= 234)^ = 22.5, p < 0.001). Marker 48 had a significantly lower OATLD prevalence than both Molasses (p < 0.001) and Carysfort (p = 0.001).

#### Amoxicillin effectiveness

Of the 66 amoxicillin-treated OATLD lesions assessed, 33% halted at the treatment line (Figure 3E). A series of generalized linear models were fitted to assess the effects of time (2019-2020 vs 2022-2023) and site, as well as their interaction, on the effectiveness of the amoxicillin treatment. The model that included the interaction effect identified no effect of site (p = 0.5), significantly higher efficacy of 2019-2020 treatments than the 2022-2023 treatments (χ² = 4.73, df = 1, p = 0.03), and a significant interaction effect between site and time period (p = 0.04). The best GLM model included only time (AIC = 84.1).

#### Seasonal assessments

The proportion of tagged *O. faveolata* colonies exhibiting OATLD was assessed for each monitoring period (2019 – 2023; Figure 4). There was a significant seasonality in the number of corals affected, peaking in October when summed across all sites (Rayleigh test of uniformity: r = 0.27, p < 0.0001). Peak prevalence varied slightly by site, with a non-significant peak in September at Sombrero (p = 0.28), and significant peaks in October at Sand Key (p < 0.005) and November at Looe Key (p < 0.0001). These seasonal outbreaks also varied by year, with the highest prevalence occurring at Sand Key in 2021, Looe Key in 2021 and 2022, and Sombrero in 2020. Though seasonal peaks were focused during annual periods of highest cumulative thermal stress, the presence of OATLD essentially disappeared during the extreme bleaching event in the summer of 2023.

**Fig. 4:**
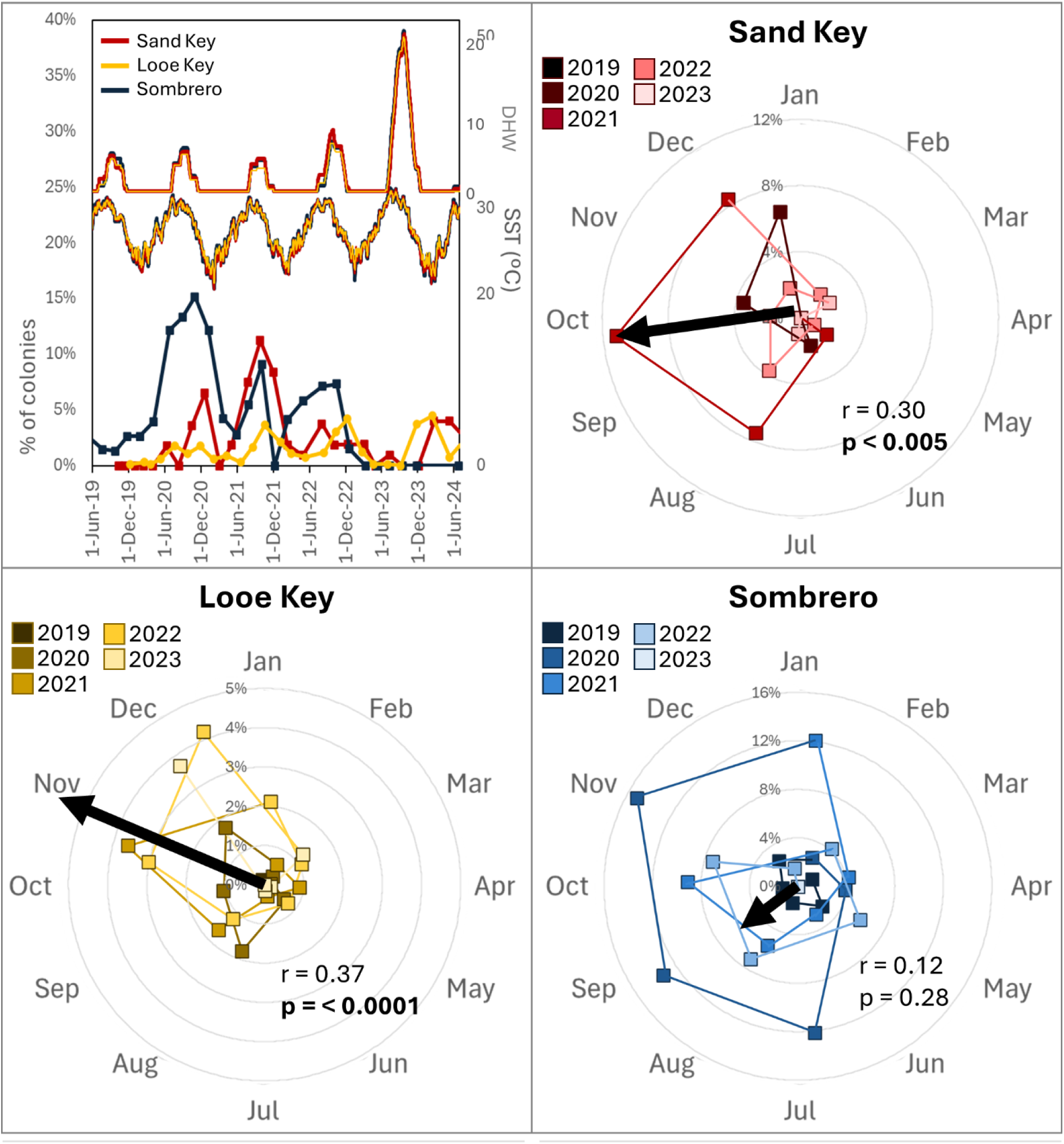
Seasonality of OATLD. The proportion of known disease-susceptible *Orbicella faveolata* colonies at three different sites exhibiting OATLD-style lesions during each monitoring event from 2019 – 2023. Sea surface temperature and degree heating weeks (DHW) data provided by NOAA Coral Reef Watch single pixel stations. Arrows represent predominant vectors (scaled to intensity) and probability of differing from random distribution (significant values in bold).

#### Colony Size

*Orbicella faveolata* colonies that were affected with OATLD at least once between 2019 and 2023 were significantly larger than colonies recorded only with SCTLD (Figure 3F). These size differences were significant at Sand Key (Wilcoxon-Rank: p < 0.001) and Looe Key (p < 0.001), but not at Sombrero. Across all three sites, OATLD-affected colonies averaged 189 ± 6 SE cm in maximum diameter, while non-OATLD colonies averaged 158 ± 3 SE cm (p < 0.001).

#### Detection of the VcpA metalloprotease

All five OATLD lesions tested negative for the VcpA metalloprotease, while four of the five SCTLD lesions also tested negative. The remaining SCTLD lesion had an irregular result, and we could not determine presence/absence. There was no significant difference in prevalence between the two groups (Fisher’s exact: p = 1 regardless of whether the erroneous result was included as positive, negative, or removed from the analysis). This suggests that *V. coralliilyticus* was not associated with the tested OATLD or SCTLD lesions.

### Microbiome Assessments

#### Sequencing

A total of 77 coral mucus-tissue slurry samples were successfully sequenced, as well as two reagent blanks, resulting in 1,608,403 raw sequencing reads. The successful samples included 28 samples of healthy tissue from healthy colonies (H), 25 unaffected tissue samples from diseased colonies (U), and 24 active disease lesion samples from those same colonies (D). The reagent blanks contained 86 reads, which were assigned to a total of 23 Amplicon Sequence Variants (ASVs). Twelve of the ASVs in the reagent blanks were identified as true contaminants with the decontam package, and these ASVs were removed from the biological samples before further analysis (Supplemental Table 2). After quality filtering and removal of contaminants, mitochondria, and chloroplasts, the dataset included 812,166 reads, with an average of 10,548 reads per sample (range: 566 – 106,879). The dataset contained 5,514 unique ASVs, which was reduced to 1,202 ASVs after removal of rare sequences that were detected in fewer than two samples or were less than 0.01% relative abundance.

#### Differential Abundance

Variation in microbial community composition was explained by the health condition of the coral sample (PERMANOVA R^2^ = 0.045, p < 0.05), but not by site (Figure 5A). Post-hoc pairwise PERMANOVA analyses showed that microbiomes from disease lesions (D) were significantly different from microbiomes of healthy tissue (H) on healthy coral colonies (p < 0.05). In addition, diseased *O. faveolata* microbiomes were more variable compared to healthy *O. faveolata* microbiomes on healthy colonies (KW: p < 0.05) (Figure 5B). However, microbiome dispersion was not significantly different between D and U samples, nor was it different between H and U samples. None of the alpha diversity metrics (richness, Chao1, and Shannon diversity) were significantly different across health conditions (Supplemental Figure 3).

**Fig. 5.**
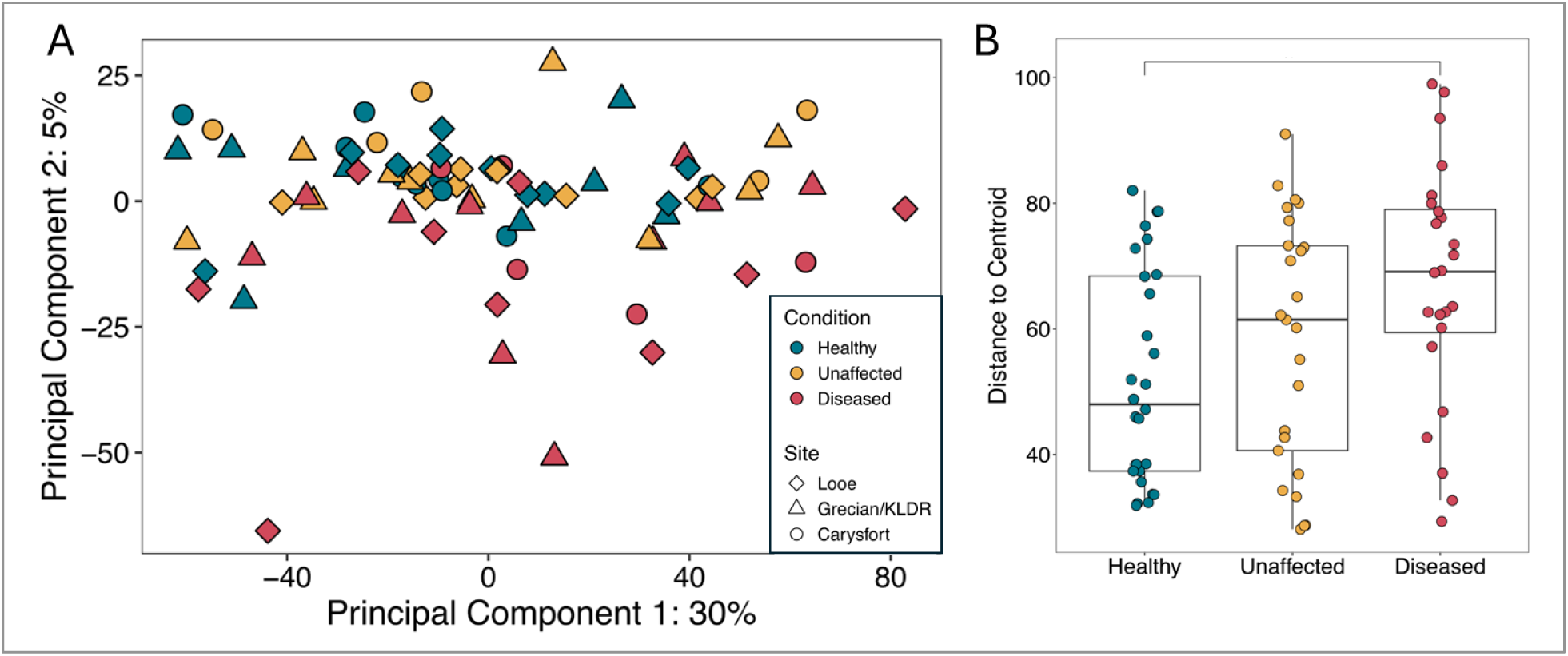
Analysis of beta diversity in microbiomes from *Orbicella faveolata* coral colonies. Microbiome samples were sourced from apparently healthy colonies, unaffected areas of diseased corals, and disease lesions on the same diseased corals. **(A)** Principal Components Analysis ordination of centered-log-ratio transformed read counts. **(B)** Beta diversity dispersion (measured as distance to centroid) across health conditions. Bracket denotes a statistically significant difference.

After determining that microbial composition differed between OATLD lesions and healthy coral colonies, we further explored how the communities changed in these two extremes. First, we used the multinomial species classification method to categorize ASVs as *O. faveolata* microbiome generalists, healthy tissue specialists, and disease lesion specialists (Figure 6A). Most ASVs were classified as either specialists for healthy coral colonies (47%) or as generalists (26%), while only 14% of ASVs were classified as disease specialists.

**Figure 6.**
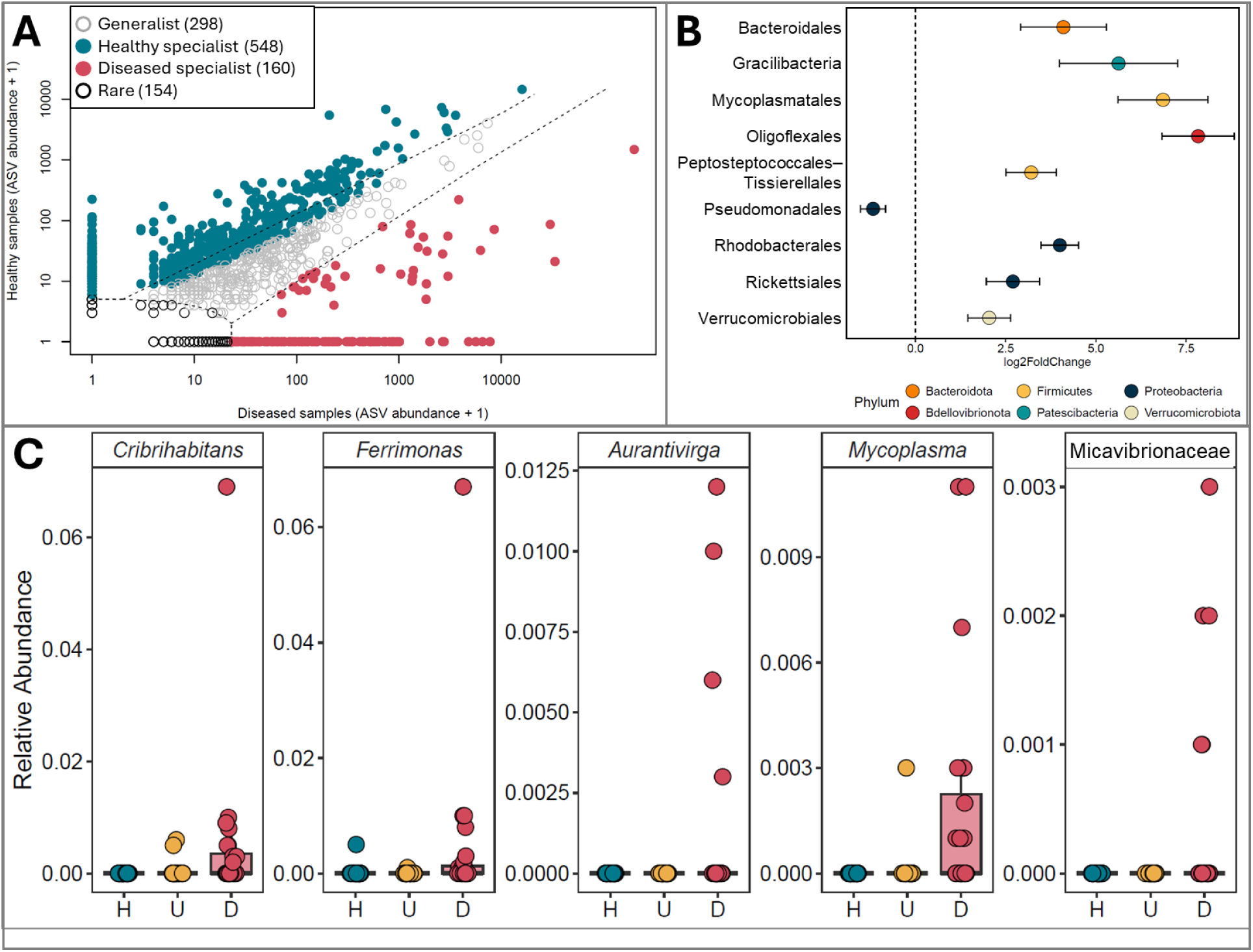
**(A)** Classification of Amplicon Sequence Variants (ASVs) as habitat specialists and generalists in microbiomes from *Orbicella faveolata* coral colonies. Only ASVs from apparently healthy (H) colonies (N = 28) or disease (D) lesions (N = 24) were included in this analysis. **(B)** Bacterial orders that were differentially abundant in microbiomes between H and D samples. Orders with positive log_2_ fold changes are enriched in disease and those with negative changes are enriched in healthy tissue. Bars represent one standard error. **(C)** Relative abundance of the five OATLD-specific ASVs that were enriched in disease lesions. Samples were sourced from healthy mucus from healthy corals (H), healthy mucus from diseased corals (U), and mucus from disease lesions (D) Lower and upper hinges on the box represent the 25th and 75th quartiles, respectively. The bars represent 1.5 times the interquartile range (between the 25th and 75th quartiles).

Next, we identified ten bacterial orders that were differentially abundant between OATLD lesions and microbiomes from healthy colonies. Nine of the ten differentially abundant bacterial orders were enriched in OATLD lesions. Only the order Pseudomonadales (class Gammaproteobacteria), comprised of 116 ASVs, was more abundant in H samples (Figure 6B). Several orders that were enriched in disease lesions were less diverse. For example, the Bacteroidales included only nine ASVs, the unclassified order of Gracilibacteria included only two ASVs, the Mycoplasmatales included only one ASV, and the Oligoflexales included eight ASVs.

Finally, we identified eighteen ASVs that were enriched in OATLD disease lesions by differential abundance analysis and were also considered to be bioindicators of OATLD based on the indicator species analysis. To pinpoint microbes specific to OATLD, we then subtracted ASVs that matched published SCTLD bioindicator sequences (Meyer et al., 2019; Becker et al., 2022) as well as ASVs belonging to genera enriched in SCTLD lesions according to the meta-analysis conducted by Rosales et al. (2023). The five remaining OATLD-specific sequences were *Cribrihabitans* ASV21, *Ferrimonas* ASV64, *Aurantivirga* ASV123, *Mycoplasma* ASV80, and Micavibrionaceae ASV158 (Figure 6C, Supplemental Table 3).

*Cribrihabitans* ASV21 (order Rhodobacterales) was an exact sequence match to 14 sequences in GenBank. Three of these sequence matches were from investigations on black band disease (GU471999, GQ455310) and associated cyanobacterial patches in corals (GQ204807) (Sato et al., 2009).The other two were from studies focused on white plague disease (KC527503, FJ203199) (Sunagawa et al., 2009; Roder et al., 2013). *Cribrihabitans* ASV21 had an average relative abundance of 0.48% in D samples, compared to 0.4% in U samples and 0% in H samples. *Cribrihabitans* ASV21 was found in nine out of 24 D samples, and two out of 25 U samples. This ASV was not detected in healthy *O. faveolata* colonies.

*Ferrimonas* ASV64 (order Enterobacterales) was an exact sequence match to 12 sequences in GenBank: FJ202753, FJ202156, FJ202638, FJ202834, FJ202813, FJ202382, FJ202123, FJ202687, FJ202195, FJ202530, FJ202858, FJ202345. All of the sequence matches for this ASV were sourced from a study on white plague disease in *O. faveolata* (Sunagawa et al., 2009). *Ferrimonas* ASV64 had an average relative abundance of 0.43% in D samples, while U and H samples had average relative abundances of 0.004% and 0.02% of the microbial community, respectively. *Ferrimonas* ASV64 was detected in nine out of 24 D samples, while it was only detected in one out of 25 U samples and one out of 28 H samples.

*Aurantivirga* ASV123 (order Flavobacteriales) was an exact sequence match for at least 100 sequences in GenBank (BLASTn hits are limited to 100 matches). The sequence matches for this ASV are ecologically diverse and have been detected in a variety of marine environments including seawater (27 sequences) and sediments (18 sequences). One sequence match was associated with the coral *Acropora millepora* (MW828552), and one sequence match was sourced from a crab with shell disease syndrome (MH061252). *Aurantivirga* ASV123 was found in 6 out of 24 D samples, with a relative abundance of 0.13%. It was not found in any U or H samples.

*Mycoplasma* ASV80 (order Mycoplasmatales) was an exact sequence match for an uncultured bacterium sequenced in the same study of white plague-diseased *O. faveolata* (FJ203048) (Sunagawa et al., 2009). *Mycoplasma* ASV80 had an average relative abundance of 0.18% in D samples, compared to 0.01% and 0% in U and H samples, respectively. *Mycoplasma* ASV80 was detected in 13 out of 24 D samples. In contrast, it was only detected in one U sample and was not detected in any of the H samples.

Micavibrionaceae ASV158 belonged to an unclassified genus (order Micavibrionales). It had no identical sequence matches within GenBank. The closest sequence match was 98.8% similar over 100% of the ASV sequence length and was an uncultured bacterium sourced from a deep-sea octocoral (DQ395935). Micavibrionaceae ASV158 had an average relative abundance of 0.04% in the D samples. It was detected in six out of 24 D sample microbiomes, but not in H or U tissues.

### Histology Assessments

Across all assessed histological parameters, there were no significant differences by reef site (Supplemental Table 4). Assessments from all sites were thus combined for sample sizes of H = 9, U = 15, and D = 15. Tabulated details of observations can be found in Supplemental Table 4.

#### Gross observations and pathology of the core samples

We observed the protrusion of mesenterial filaments (MFs; Figure 7) in 100% of healthy (H) samples and 87% of U samples, with no significant difference between these groups (Figure 10; Fisher’s: p = 0.51). In contrast, only 27% of D samples had protruding MFs, significantly less than in both H and U samples (p < 0.003). The protruded MFs appeared white and/or orange as they extended from pores or gaps in the coenenchyme and the mouths of the polyps.

**Figure 7.**
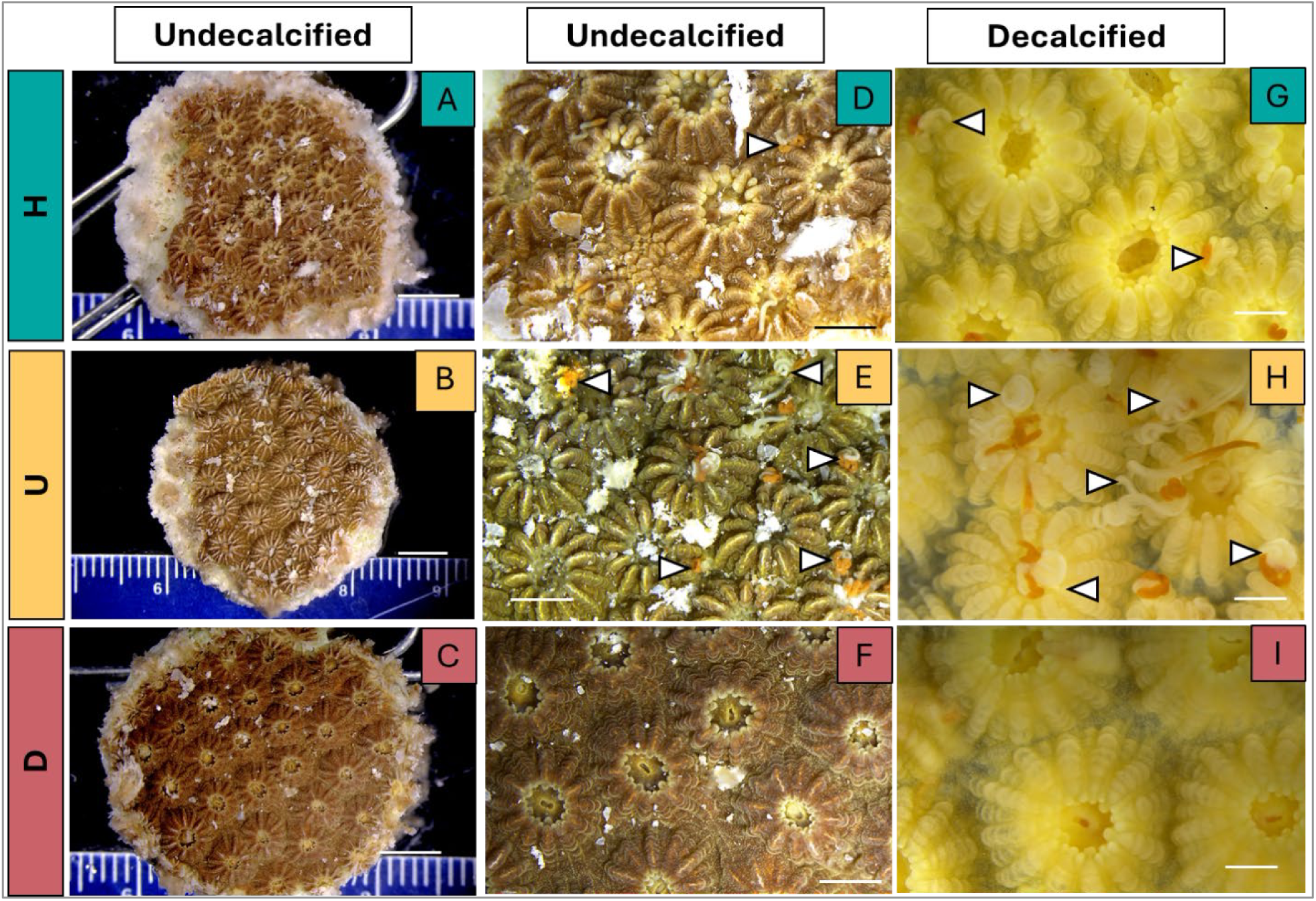
Macrophotographs of fixed *Orbicella faveolata* samples from a healthy (“H”) colony (panels **A,D,G**), unaffected (“U”) tissue on a diseased colony (panels **B, E, H**), and adjacent to an OATLD disease (“D”) lesion (panels **C, F, I**) High magnification of undecalcified views (D, E, F) show a light mesenterial filament (MF) protrusion in D, a moderate amount in E, and no MF protrusion in F. Decalcified views (G, H, I) again show a light amount of MF protrusion in G, a moderate amount in H, and none in I. Arrowheads indicate MFs. Scale bars: 5mm (A–C), 2mm (D – F), and 1mm (G – I).

#### Histopathology

We primarily observed surface body wall lesions in the gastrodermis, predominantly affecting the endosymbionts. Necrotic MFs and associated mesenteries were sometimes found protruding through the surface body wall (Figure 8A) or contained entirely within the gastrovascular cavities (Figure 8B). Otherwise, we found no prominent pathological changes of the basal body wall gastrodermis. We noted histopathological cellular changes and responses of the host tissue in the MFs, particularly in the density and appearance of the granular amoebocytes, mucus cell (mucocyte) hypertrophy, and pathological changes within the endosymbionts.

**Figure 8.**
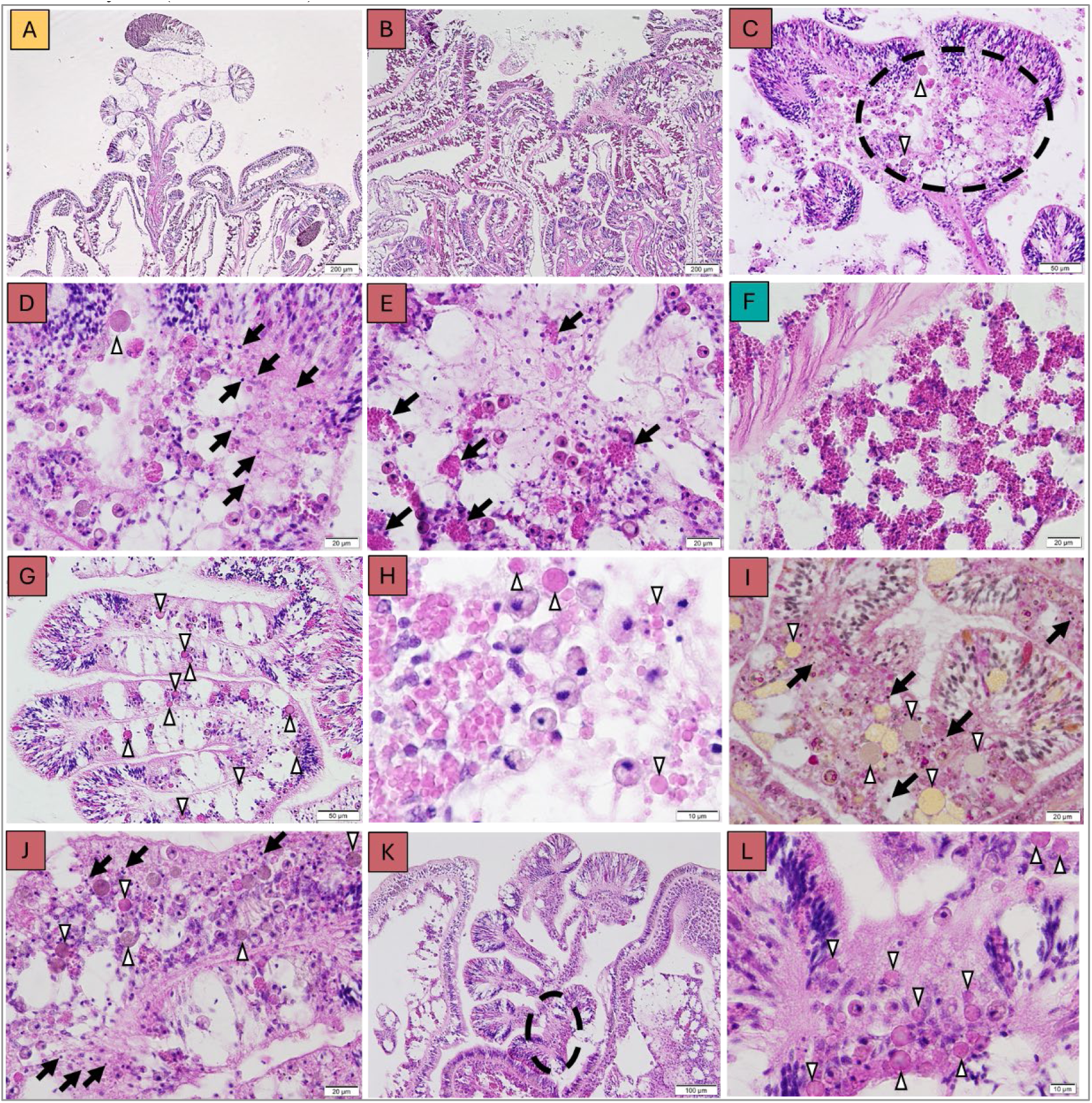
Histological sections (H&E stain, except for (I) which is PAS-MY) within the mesenterial filaments (MFs) of *Orbicella faveolata*. All sections are from diseased samples except A (unaffected tissue from a diseased colony) and F (healthy). **(A)** MFs protruding through the surface body wall in unaffected tissue from a diseased colony. **(B)** MFs (cnidoglandular bands and associated mesenteries) fill the aboral gastrovascular cavities (lower half of image). **(C)** Necrosis (dotted circle) of MFs associated with degraded granular amoebocytes (arrowheads). **(D)** High magnification of dotted circle area from (C) showing karyorrhexis (arrows) and degraded granular amoebocytes (arrowheads). **(E)** Depletion (aggregation) of granular amoebocytes (arrows). **(F)** Abundant, intact granular amoebocytes from an apparently healthy colony. **(G)** Degraded granular amoebocytes (globules; arrowheads) in the MFs. **(H)** High magnification of degraded granular amoebocytes (arrowheads). **(I, J)** Karyorrhexis (arrows) and degraded granular amoebocytes (arrowheads). **(K)** Protruded MFs. **(L)** High magnification of the dotted circled area from (K; rotated 90°) showing degraded granular amoebocytes (arrowheads).

#### Necrotic gastrodermis

We did not observe lytic necrosis of the gastrodermis as described in the SCTLD samples of Landsberg et al. (2020). However, at the presumed tissue-loss margin of a single D sample, the surface body wall gastrodermis did exhibit karyorrhexis (Supplemental Figure 1).

#### Necrotic mesenterial filaments (MFs)

Necrotic MFs (primarily within the polyps’ gastrovascular cavities; Figure 8C-D) were significantly more common (Fisher’s: p < 0.001) in D samples (87%), than in U (13%) or H (0%) samples, with no significant difference in prevalence between U and H samples (Figure 10).

#### Depleted (aggregated) granular amoebocytes

The loss of granular amoebocytes due to the cell aggregation (Figure 8E) in the mesenteries and associated MFs (mainly in the gastrovascular cavities) was significantly more frequent in D samples (100%; Fisher’s: p < 0.001) than in the U (27%) and H (22%) samples, with no significant differences between U and H (Figure 10). Apparently healthy samples usually contained abundant granular amoebocytes in the MFs (Figure 8F).

#### Degraded granular amoebocytes and associated globules

We found degraded granular amoebocytes in the MFs associated with cnidoglandular bands (Figure 8G-J). These were significantly more common in D samples (93%; Fisher’s: p < 0.002) than in U (33%) and H (0%) samples; the proportion of U and H samples that contained degraded granular amoebocytes did not differ significantly (Figure 10).

Rounded spherical globules, approximately 10 µm in diameter (range 7.5–12.5 µm) and with a homogeneously flat to bumpy appearance, were found in the mesenteries associated with MFs throughout the gastrovascular cavities of the polyps. They were also detected, but rarely, in the externally protruded MFs, notably in D samples (Figure 8K, L). The globules were presumably associated with necrosis of granular amoebocytes in the MFs (Figure 8D, I, J). They stained eosinophilic to brownish with H&E (Figure 8D, H, J, L), light yellow to brownish with PAS-MY (Figure 8I), and occasionally possessed a thickened eosinophilic outer ring.

#### Mucocyte hypertrophy

Mucocytes in the MFs (mainly within the gastrovascular cavities) appeared to be enlarged. They contained abundant mucus that stained prominently purple to blue with thionin (Figure 9A–D). We also observed extraneous mucus near these hypertrophic mucocytes. Within the adjacent cnidoglandular band, we found dissociated granules containing mucin. Mucocyte hypertrophy was confirmed across all health conditions (H = 11%, U = 33%, D = 47%), with no significant differences among them.

**Figure 9.**
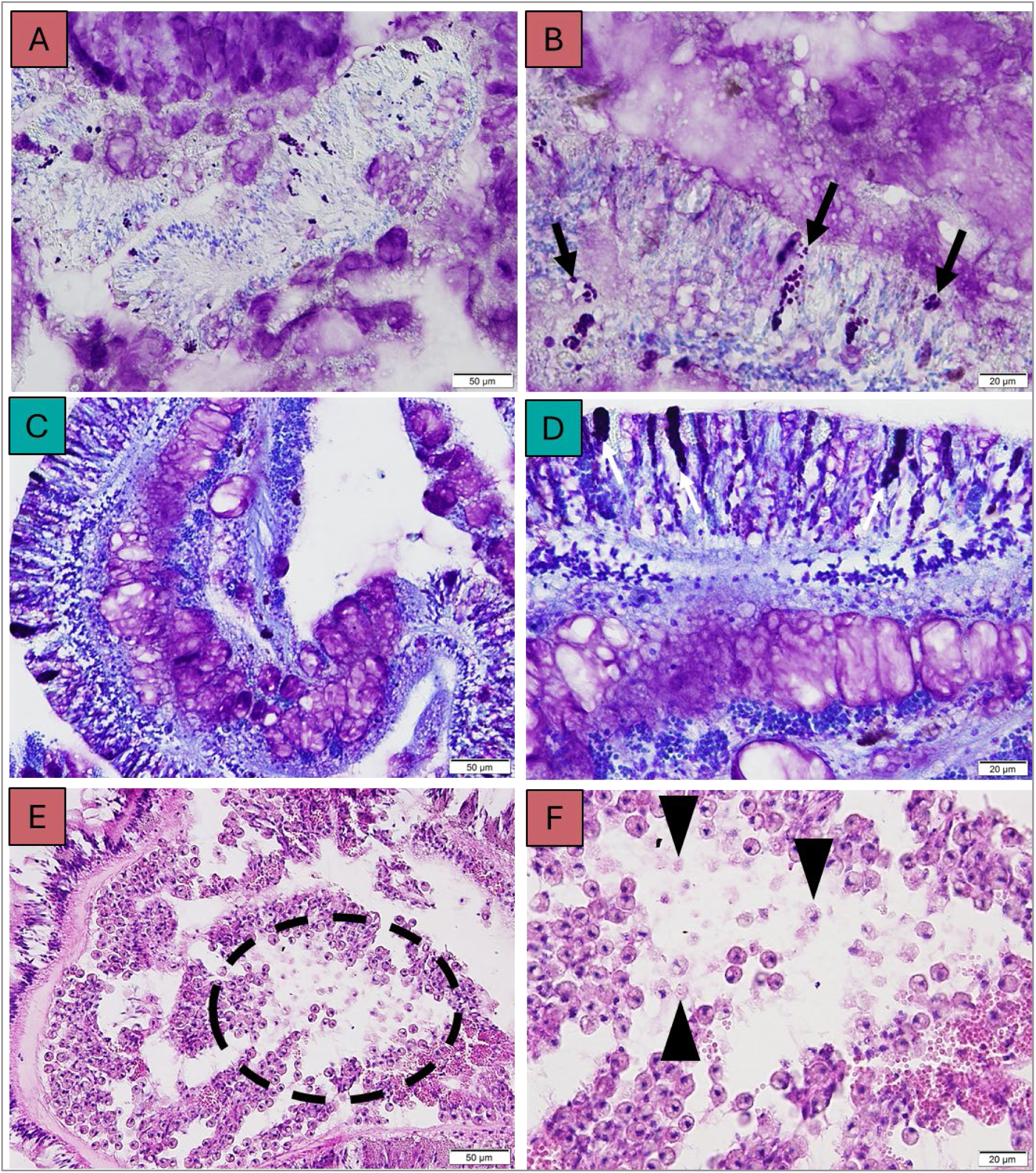
Histological sections of *Orbicella faveolata* in the mesenterial filaments (MFs; **A–D**: thionin) and surface body wall of the gastrodermis (**E,F**: H&E stain). **(A)** Disease-affected (D) samples with copious amounts of mucus and mucocyte hypertrophy. **(B)** is a high magnification section of (A). Mucus stained light purple - brown, and granules containing mucin in the cnidoglandular band adjacent to the MFs with hypertrophied mucocytes were dissociated (arrows). (**C**) Healthy (H) samples with normal mucus production in the MF. **(D)** is a high magnification section of (C). Mucus stained dark purple, with a condensed appearance, and the granules were tightly condensed (arrows). (**E**) Focal necrosis of endosymbionts (dotted circle) in the surface body walls of the gastrodermis of a disease-affected sample. (**F**) High magnification views of the circle in (E) showing necrotic endosymbionts (arrowheads).

#### Periodic acid–Schiff (PAS)-positive reaction of endosymbionts

Starch granules in the cytoplasm of the endosymbionts located in the surface body wall gastrodermis stained strongly positive (red) with PAS-MY (Supplemental Figure 2) in all specimens except one H sample.

#### Necrosis of endosymbiont (ghosting)

“Ghosting” of an endosymbiont is the result of multifocal necrosis which retains only the outer cell wall frame of the remnant endosymbiont body, visible as a pale stain with H&E. Internal structures of the nucleus and other organelles undergo apparent loss (Figure 9E, F). We observed endosymbiont ghosting in the surface body wall gastrodermis. These endosymbionts exhibited somewhat swollen, light eosinophilic, or occasionally brownish-green tinted cytoplasm when stained with H&E. Degenerative changes occurred, nuclei were absent, and an enlarged symbiosome space had presumably expanded and coalesced with adjacent symbiosomes. Endosymbiont ghosting was significantly more frequent in D (93%) than U (40%) and H (11%) (Figure 10; Fisher’s: p < 0.0001), and there was no statistical difference between U and H (p = 0.19).

**Figure 10.**
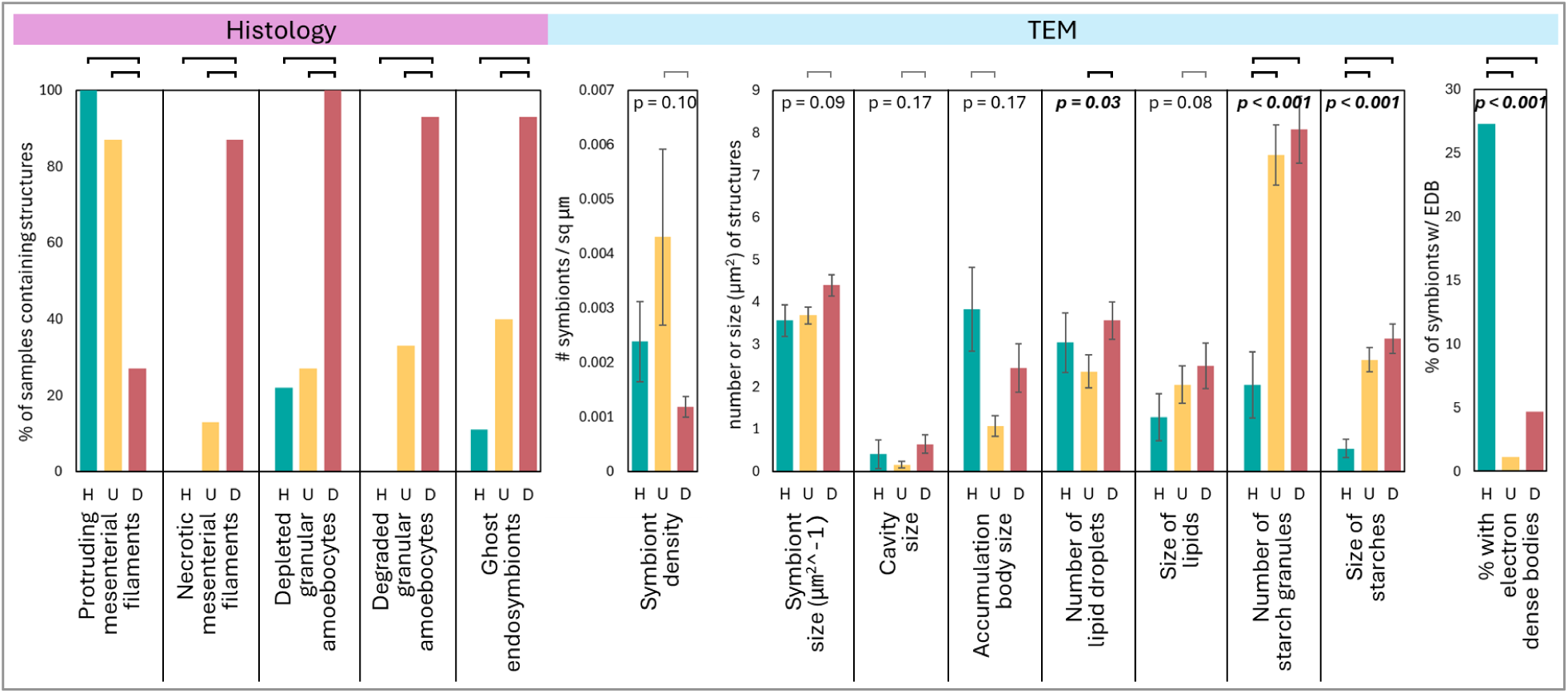
Prevalence of histological parameters (including protrusion of mesenterial filaments at the surface in post-fixed core samples by dissecting microscope) and physiological metrics of TEM in *Orbicella faveolata* samples from healthy (H) colonies, unaffected areas of diseased corals (U), and disease-adjacent tissues (D). Error bars show standard error where applicable. Bracket lines over the bars show significant differences between health groups. For histological parameters, Fisher’s exact tests with an α < 0.05 are considered significant. Numbers above the TEM assessment bars indicate the Kruskal-Wallis p-values, with significant differences in bold italics. Significant post-hoc Dunn tests are shown as brackets for all TEM variables, but are faded for metrics where the Kruskal-Wallis values were not significant.

#### Organisms

In 7 of the samples (4 from Looe Key and 3 from Carysfort), apicomplexan sporozoites (Landsberg et al., 2020; Hawthorn et al., 2024) were occasionally detected in the lobes of the MFs regardless of health condition. Endolithic algae-hyphae were only sporadically and lightly detected in the samples. Possible coccoid-like, coccobacilloid-like structures (Landsberg et al., 2020) that stained magenta with Giemsa were observed frequently in the surface body wall gastrodermis.

### TEM Assessments

#### Density and size of symbionts

Symbiont density did not differ significantly when all health state groups were compared (KW: p = 0.1). However, diseased samples (D) averaged 0.001 symbionts / μm^2^ of gastrodermis tissue, which was significantly lower than symbiont density of unaffected tissue (U) from diseased corals (0.004 symbionts / μm^2^; Dunn: p = 0.02). Samples from healthy corals (H) had an intermediate density (0.002 symbionts / μm^2^), which was not significantly different from the other groups (Figure 10).

Symbiont size did not differ when all health states were compared (KW: p = 0.09). However, in pairwise comparisons, symbionts of D samples were significantly larger (44 μm^2^ ± 2.5 SE) than those of U samples (37 μm^2^ ± 2 SE. Dunn: p = 0.03). Symbiont size did not differ between the D and H (36 μm^2^ ± 4 SE) samples, likely due to the small number of H samples (Figure 10).

#### Symbiont degradation

Across the 174 symbionts scored, only 24 exhibited degradation. Symbiont degradation was observed in 23% of H samples, 16% of U samples, and 8% of D samples; there was no significant difference among health states.

#### Cavity presence and size

The presence of cavities within symbionts (Figure 11C) did not differ significantly among health states (H = 9%, U = 8%, D = 17%). The size of the cavities also did not differ among all health states (KW: p = 0.17), but in pairwise tests, symbionts from D corals did have larger cavities (0.6 μm^2^ ± 0.2 SE) than those in U symbionts (0.2 μm^2^ ± 0.1 SE; Dunn: p = 0.03; Figure 10). This trend for differences in cavity size among health states needs to be further explored with a larger sample size, as cavity size was highly variable and there were only a small number of samples containing cavities (H = 2, U = 7, D = 11).

**Figure 11.**
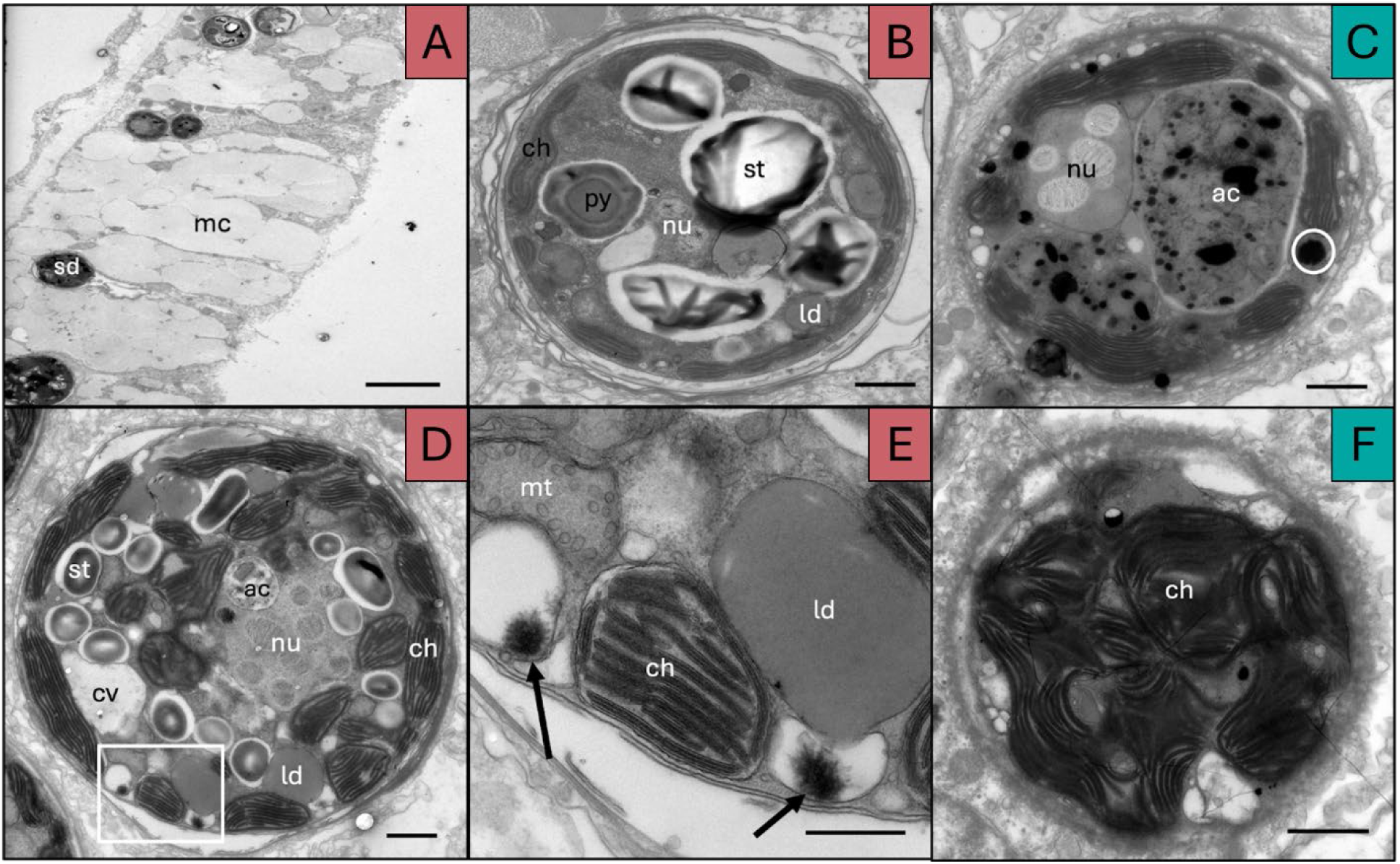
Transmission electron micrographs of *Orbicella faveolata* tissue and Symbiodiniaceae within the tissue. **(A)** An overview of the gastrodermis from a diseased coral sample, highlighting the symbionts and mucocytes. **(B)** A Symbiodiniaceae within diseased coral tissue showing starch granules, lipid droplets, intact chloroplasts, a pyrenoid with a starch sheath, and a nucleus. **(C)** A Symbiodiniaceae within a healthy coral with accumulation bodies and electron dense bodies (white circle). **(D)** A Symbiodiniaceae in a diseased coral with starch granules, viral-like particles (VLPs), an accumulation body, a cavity, lipid droplets, a nucleus, and intact chloroplasts, (**E)** A zoomed in micrograph of the white box in (D) with detailed stellate VLP structures (black arrows), thylakoid membranes within the chloroplasts, mitochondria, and a lipid droplet. **(F)** A degraded Symbiodiniaceae within a healthy coral exhibiting chloroplasts but no other organelles present. Structures are labeled: sd = Symbiodiniaceae, mc = mucocyte, ch = chloroplast, st = starch granule, py = pyrenoid, ld = lipid droplet, nu = nucleus, ac = accumulation body, mt = mitochondria, cv = cavity. Scale bars are: 10 µm for (A), 1 µm for (B, C, D, and F), and 500 nm for (E).

#### Accumulation body presence and size

Within the symbionts, accumulation bodies were found in 50% of H samples, 25% of U samples, and 28% of D samples. There were no significant differences among groups. The size of the accumulation bodies also did not differ across all groups (Kruskal-Wallis: p = 0.17). However, post-hoc tests identified H samples as having larger bodies (3.8 μm^2^ ± 1.0 SE) than U samples (1.1 μm^2^ ± 0.2 SE; Dunn: p = 0.003). Accumulation body size was moderately correlated with the symbiont size (Spearman correlation coefficient = 0.63) for H samples but showed no significant correlations for U or D samples.

#### Presence of degraded chloroplasts, degraded thylakoids, degraded nuclei, and membrane separation from symbiosome

Chloroplast degradation was seen in 45% of H samples, 48% of U samples, and 31% of D samples. We observed thylakoid degradation in 41% of H samples, 43% of U samples, and 30% of D samples. Of the 68 symbionts with a visible nucleus, we documented nucleus degradation in 23% of H samples, 25% of U samples, and 34% of D samples. Across all samples, 28 cells had membrane separation from the symbiosome. This included 14% of H samples, 19% of U samples, and 13% of D samples. There were no significant differences among health states for any of these parameters.

#### Number and size of lipid droplets within symbionts

The number of lipid droplets (Figure 11) differed significantly among health states (KW: p = 0.03). In pairwise comparisons, symbionts from D samples contained significantly more (3.6 ± 0.4 SE) lipid droplets than symbionts from U samples (2.4 ± 0.4 SE; Dunn: p = 0.004). The number of lipid droplets within symbionts from H samples (3.0 ± 0.7 SE) did not significantly differ from either D or U (Figure 10).

The size of the lipid droplets did not differ significantly among all groups (KW: p = 0.08). In pairwise comparisons, symbionts from D corals had a larger lipid area (2.5 μm^2^ ± 0.5 SE) than U corals (2.1 μm^2^ ± 0.4 SE; Dunn: p = 0.02). Healthy corals had the smallest lipid area (1.3 μm^2^ ± 0.6 SE), but this was not significantly different from U or D groups (Figure 10).

#### Number and size of starch granules within symbionts

Overall, the number of starch granules (Figure 11) within D and U samples were greater than those from H samples (KW: p < 0.001; Figure 10). Symbionts within H samples contained 2.0 ± 0.8 SE starch granules, significantly less than the number in U samples (7.5 ± 0.7 SE; Dunn: p < 0.0001) and D samples (8.1 ± 0.8 SE; p < 0.0001).

The starch reserves within D and U samples had a greater area than those of H samples (KW: p < 0.001; Figure 10). In H samples, starch reserve area averaged 0.5 μm^2^ ± 0.2 SE, significantly less than those in U samples (2.6 μm^2^ ± 0.3 SE; Dunn: p < 0.0001) and D samples (3.1 μm^2^ ± 0.4 SE; Dunn: p < 0.0001).

Symbionts in D samples had a strong correlation between symbiont size and number of starch granules (Spearman correlation coefficient = 0.75) as well as starch area (Spearman correlation coefficient = 0.83). Symbionts from U samples also exhibited strong correlations to the number of starch granules (Spearman correlation coefficient = 0.80) and starch area (Spearman correlation coefficient = 0.76). There were no significant correlations between symbiont size and starch area for H samples.

#### Pyrenoid and starch sheath

Forty eight of the assessed symbionts contained a pyrenoid (Figure 11B). There were no significant differences between health states (H = 45%, U = 27%, D = 22%). All of the symbionts containing a pyrenoid also had a starch sheath surrounding it, and did not differ significantly between health states. Notably, all corals were collected and preserved during the day.

#### VLP presence

Viral-like particles (VLPs; Figure 11D) were consistently observed across all health states, with no significant differences among groups. Seventy seven percent of H symbionts, 68% of U symbionts, and 75% of D symbionts contained VLPs. A single viroplasm was found within one symbiont in a D sample.

#### Electron dense body presence

Electron dense bodies (Figure 11C) were found in 10 samples, including 27% of H samples, 1% of U samples, and 5% of D samples. The presence of these bodies was significantly higher in H samples than in U samples (Fisher: p < 0.001) and D samples (Fisher: p = 0.02; Figure 10).

#### Comparison to SCTLD from Work et al. 2021

Within the Work et al. (2021) SCTLD dataset, cavities within endosymbionts were found in both H and D samples (Figure 12A); all three health states of the OATLD samples also contained cavities. There was a significant difference between diseases and health state samples (KW: p = 0.04); OATLD U samples (6.2% ± 2.6 SE) had a lower prevalence of cavities than both SCTLD H (37% ± 0 SE; Dunn: p = 0.01) and SCTLD D (46.5% ± 6.5 SE; Dunn: p = 0.003) samples.

**Figure 12.**
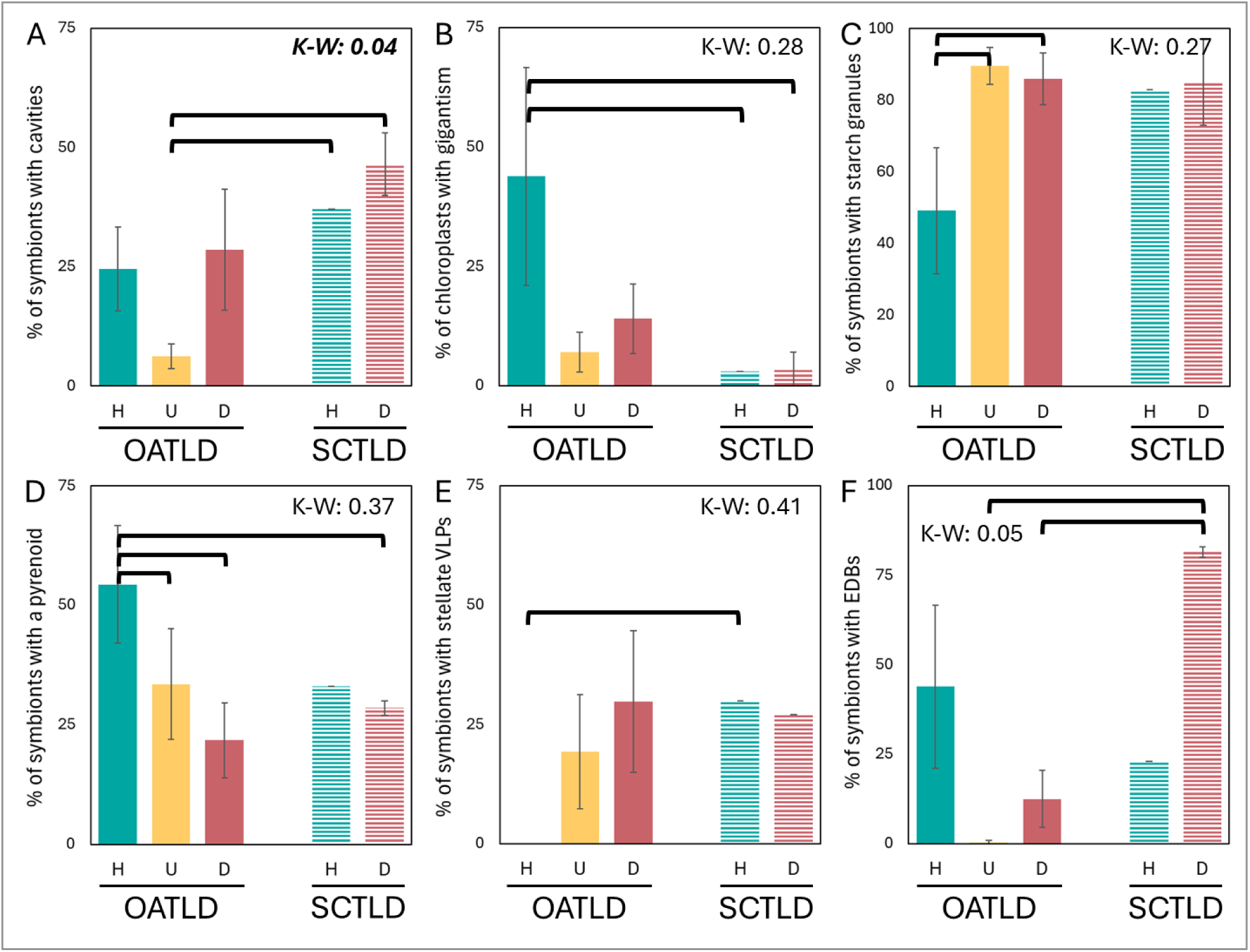
Comparison of physiological metrics of *Orbicella faveolata* symbionts across two diseases (OATLD and SCTLD) and health states (H = healthy, U = unaffected tissues of diseased colonies, D = diseased tissues). **(A)** The percentage of symbionts containing cavities, **(B)** the percentage of symbionts with chloroplasts exhibiting gigantism, **(C)** the percentage of symbionts containing starch granules, **(D)** the percentages of symbionts with a pyrenoid, **(E)** the percentage of symbionts containing stellate viral-like particles (VLPs) and **(F)** the percentage of symbionts containing electron dense bodies (EDBs). Error bars indicate standard error. K-W numbers indicate the p-value for Kruskal-Wallis tests across the disease/health states; significant values (panel A) are in bold italics. Bracket lines above the bars indicate values that differed significantly from each other in pairwise Dunn’s comparisons, regardless of the significance of the Kruskal-Wallis test.

The prevalence of chloroplast gigantism within endosymbionts did not differ among diseases and health states (KW: p = 0.28; Figure 12B). However, in pairwise comparisons, chloroplast gigantism was more common in OATLD H samples (43.9% ± 22.8 SE) than in SCTLD H (3% ± 0 SE; Dunn: p = 0.04) and SCTLD D (3.5% ± 3.5 SE; Dunn: p = 0.03) samples.

The presence of thylakoid degradation did not differ among any of the health or disease states in either the Kruskal-Wallis or post-hoc Dunn’s comparisons.

The presence of starch granules within symbionts did not vary across health states and diseases (KW: p = 0.27; Figure 12C). However, in post-hoc tests, OATLD H samples had a lower proportion of symbionts with starch granules (49.1% ± 17.5 SE) than OATLD U (89.6% ± 5.1 SE; p = 0.02) and OATLD D (86.0% ± 7.3 SE) samples; this difference is supported by the raw data analyses for which the proportion of H symbionts containing starch granules was significantly lower than U and D samples (Figure 10). Among the *O. faveolata* SCTLD samples, there were no differences in the proportion of symbionts containing starch granules between the D and H samples. However, the larger multi-species SCTLD dataset indicated that the proportion of symbionts containing starch granules was higher in healthy corals than in SCTLD-affected ones (Work et al., 2021), which contrasts with the OATLD results.

The absence of a pyrenoid was common in the description of SCTLD (Work et al., 2021). However, we found no differences in the prevalence of symbionts containing pyrenoids across OATLD and SCTLD health states (KW: p = 0.37; Figure 12D). Follow-up pairwise comparisons did suggest that the prevalence of a pyrenoid was greater in OATLD H samples (54.4% ± 12.3 SE) compared to OATLD U (33.5% ± 11.6 SE; Dunn: p = 0.05) and OATLD D (21.8% ± 7.9 SE; Dunn: p = 0.04) samples. OATLD H samples also had a higher prevalence of pyrenoids within the symbionts compared to SCTLD D samples (28.5% ± 1.5 SE; Dunn: p = 0.04). These differences may be affected by the small sample size of OATLD H symbionts.

The presence of stellate VLPs was also a focal point in SCTLD signs by Work et al. (2021). However, among both diseases and all health states, there were no significant differences in the proportion of symbionts containing stellate VLPs (KW; p = 0.41; Figure 12E). Post-hoc pairwise comparisons identified no significant difference in the presence of stellate VLPs between SCTLD H and SCLTD D samples. However, SCTLD H samples did have a higher prevalence of stellate VLPs (30% ± 0 SE) than OATLD H samples (0% ± 0 SE; Dunn: p = 0.03).

Prevalence of electron dense bodies within the symbionts trended towards but did not significantly vary among diseases and health states (KW: p = 0.05; Figure 12F). Post-hoc pairwise comparisons suggested that symbionts from SCTLD D samples had a higher prevalence of electron dense bodies (81.5% ± 1.5 SE) than OATLD U (0.4% ± 0.4 SE; Dunn: p = 0.003) and OATLD D (12.5% ± 8.0 SE; Dunn: p = 0.02) samples. Though not significantly different, healthy OATLD corals in this analysis had a higher prevalence of electron dense bodies than the OATLD U and D samples, which is supported by the raw data comparisons (Figure 10).

## Discussion

Tissue loss lesions have been reported on multiple species of non-acroporid corals in the Caribbean since the late 1970s, identified primarily as white plague types I-III and stony coral tissue loss disease (SCTLD). White plague type I (initially termed “plague”) was identified by (Dustan, 1977) on at least seven species in the Florida Keys, including the *Orbicella* species complex as well as *Mycetophyllia* spp., *Colpophyllia natans*, *Porites astreoides*, and *Stephanocoenia intersepta*. Almost 20 years later in 1995, another outbreak in the Florida Keys was characterized by Richardson et al. (1998a). This outbreak was termed white plague type II, had faster lesion progression rates, and impacted at least 17 species, including *Orbicella* spp., but most notably affecting *Dichocoenia stokesii*. Four years later in 1999, another Florida Keys outbreak was described as white plague type III (Richardson et al., 2001). This outbreak was described as causing extremely rapid tissue loss on large corals, specifically *Colpophyllia natans* and *O. faveolata* (lumped into the *annularis* species complex in Richardson et al. (2001)). However, the white plague type III nomenclature was discarded due to inconsistent pathogen identification, with only white plague type I and type II subsequently considered valid (Cróquer et al., 2021). SCTLD emerged in southeast Florida in 2014 (Precht et al., 2016) and subsequently spread throughout the Caribbean, impacting over half of the region’s reef building coral species (Kramer et al., 2025). SCTLD impacts at least 23 species of corals (Florida Coral Disease Response Research & Epidemiology Team, 2018), often sweeping through reefs at epidemic proportions.

The etiological agent of most coral diseases, including SCTLD, remains unknown. Attempts to characterize a singular putative pathogen are often confounded by the complex symbiosis of the coral holobiont, the presence of opportunistic microbes, and our inability to successfully culture some pathogenic microbes for the fulfillment of Koch’s postulates (Vega Thurber et al., 2020). Furthermore, some coral diseases are likely the result of a consortium of microbes acting as a polymicrobial infection rather than a single pathogen causing disease (Meyer et al., 2025). The pathogen for white plague type II was identified through the fulfillment of Koch’s postulates as a *Sphingomonas* bacterium (Richardson et al., 1998b) and later established as the novel species *Aurantimonas coralicidia* (Denner et al., 2003). However, subsequent research did not detect *A. coralicidia* in *O. faveolata* with white plague type II (Sunagawa et al., 2009), and low relative abundances of *A. coralicidia*-like amplicons were detected in both healthy and diseased tissues of *O. annularis* with white plague-like symptoms (Cook et al., 2013). For white plague type I, white plague type III, and SCTLD, no pathogens have yet been identified.

Here we discuss the findings from our OATLD samples in a comparative context with SCTLD and white plague types, exploring similarities, differences, and future directions.

### Field Assessments

Assessments of OATLD in the field identified few similarities to SCTLD. In contrast, many but not all features resembled white plague.

#### SCTLD

The overall appearance and behavior of lesions contradicted that of known SCTLD lesions in many ways (Table 1). Compared to SCTLD lesions on *O. faveolata*, observed OATLD lesions were smoother (less undulating) and more linear (Figure 2). Lesions also behaved differently from those of SCTLD in that 44% of them halted on their own within two months, whereas SCTLD lesions generally consume all available tissue. Additionally, when treated with amoxicillin, only 33% of lesions halted at the treatment line. In contrast, amoxicillin treatments on *O. faveolata* SCTLD lesions have 91 to 93% efficacy on SCTLD-affected *O. faveolata* in the Florida Keys (Neely et al., 2020; Neely et al., 2021b). Seasonality of prevalence also differed between OATLD and SCTLD. While OATLD prevalence significantly increased during or immediately following peak thermal stress months, long-term monitoring of SCTLD-affected colonies at numerous offshore Florida Keys reefs from 2019 to 2023 identified a steadily declining, but non-seasonal trend in prevalence (Neely, 2023a). Overall, the appearance and behavior of these OATLD lesions does not match what is known about SCTLD on these reefs.

**Table 1:**
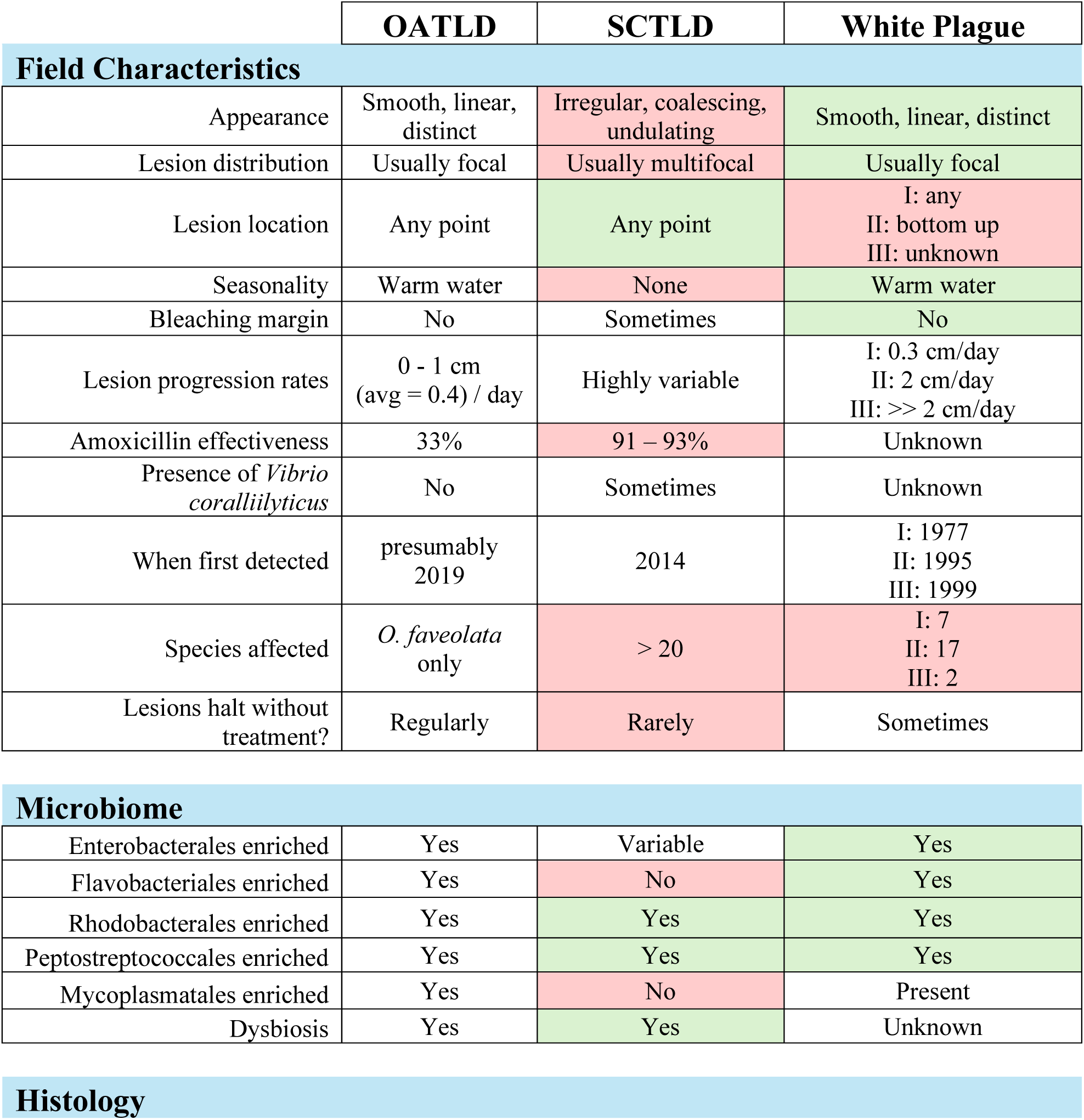
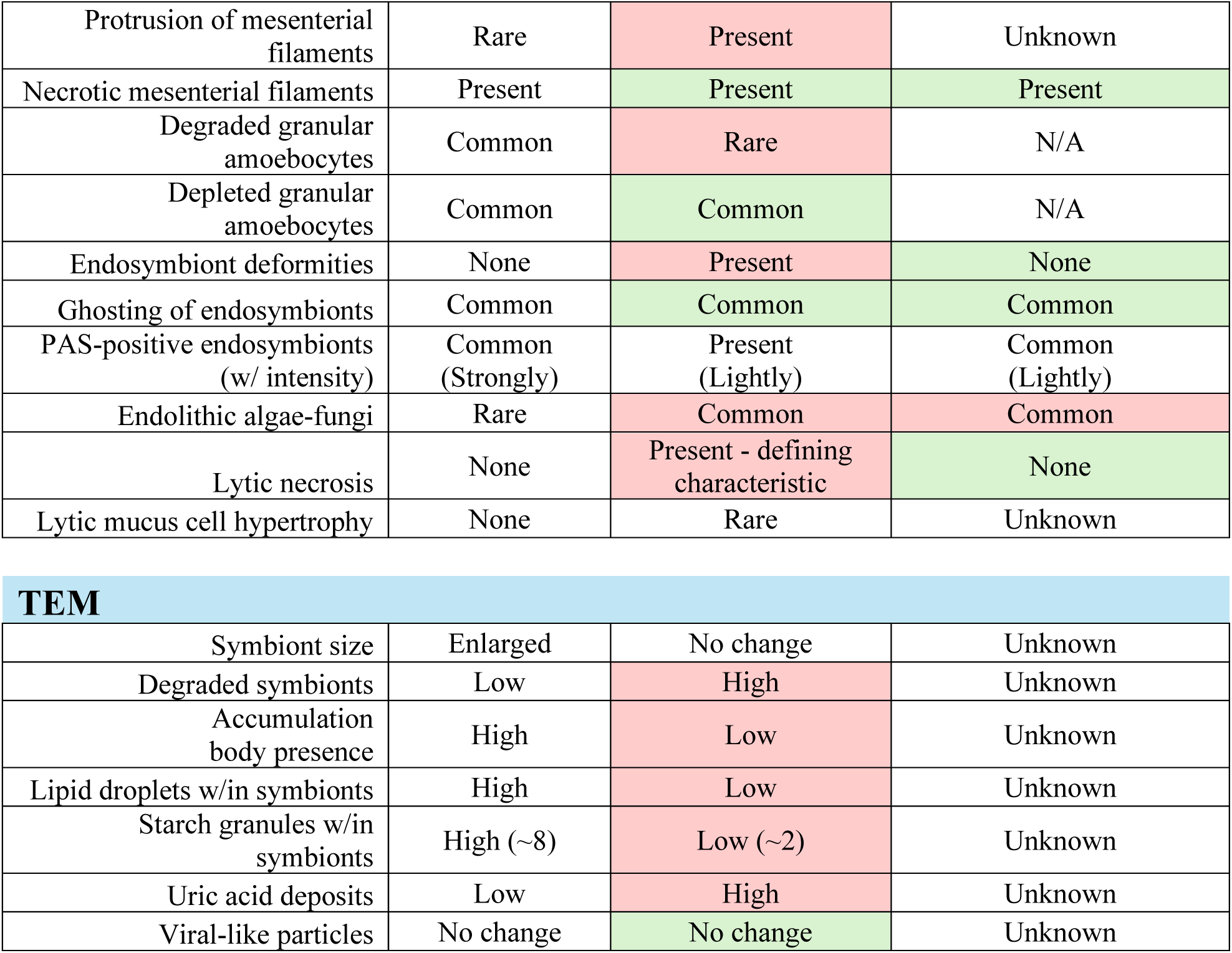
Summary of comparisons between OATLD samples and known characteristics of SCTLD and white plague syndromes. I, II, and III indicate distinctions among the three white plague types. Red cells indicate differences from OATLD, and green cells indicate similarities. The comparisons to white plague disease in the histology section are to *Orbicella franksi* specimens (E. Peters, unpublished data); *O. franksi* do not have granular amoebocytes in the MFs, so those comparisons are not applicable (N/A).

|  | OATLD | SCTLD | White Plague |
| --- | --- | --- | --- |
| **Field Characteristics** |  |  |  |
| Appearance | Smooth, linear, distinct | Irregular, coalescing, undulating | Smooth, linear, distinct |
| Lesion distribution | Usually focal | Usually multifocal | Usually focal |
| Lesion location | Any point | Any point | I: any<br>II: bottom up<br>III: unknown |
| Seasonality | Warm water | None | Warm water |
| Bleaching margin | No | Sometimes | No |
| Lesion progression rates | 0 - 1 cm<br>(avg = 0.4) / day | Highly variable | I: 0.3 cm/day<br>II: 2 cm/day<br>III: >> 2 cm/day |
| Amoxicillin effectiveness | 33% | 91 – 93% | Unknown |
| Presence of *Vibrio coralliilyticus* | No | Sometimes | Unknown |
| When first detected | presumably 2019 | 2014 | I: 1977<br>II: 1995<br>III: 1999 |
| Species affected | *O. faveolata* only | > 20 | I: 7<br>II: 17<br>III: 2 |
| Lesions halt without treatment? | Regularly | Rarely | Sometimes |
| **Microbiome** |  |  |  |
| Enterobacterales enriched | Yes | Variable | Yes |
| Flavobacteriales enriched | Yes | No | Yes |
| Rhodobacterales enriched | Yes | Yes | Yes |
| Peptostreptococcales enriched | Yes | Yes | Yes |
| Mycoplasmatales enriched | Yes | No | Present |
| Dysbiosis | Yes | Yes | Unknown |
| Histology |  |  |  |
| Protrusion of mesenterial filaments | Rare | Present | Unknown |
| Necrotic mesenterial filaments | Present | Present | Present |
| Degraded granular amoebocytes | Common | Rare | N/A |
| Depleted granular amoebocytes | Common | Common | N/A |
| Endosymbiont deformities | None | Present | None |
| Ghosting of endosymbionts | Common | Common | Common |
| PAS-positive endosymbionts (w/ intensity) | Common (Strongly) | Present (Lightly) | Common (Lightly) |
| Endolithic algae-fungi | Rare | Common | Common |
| Lytic necrosis | None | Present - defining characteristic | None |
| Lytic mucus cell hypertrophy | None | Rare | Unknown |
| TEM |  |  |  |
| Symbiont size | Enlarged | No change | Unknown |
| Degraded symbionts | Low | High | Unknown |
| Accumulation body presence | High | Low | Unknown |
| Lipid droplets w/in symbionts | High | Low | Unknown |
| Starch granules w/in symbionts | High (~8) | Low (~2) | Unknown |
| Uric acid deposits | Low | High | Unknown |
| Viral-like particles | No change | No change | Unknown |

The absence of the VcpA metalloprotease on the tested OATLD lesions (and the lack of enriched *Vibrio* relative abundance as indicated by the microbiome analyses) suggests that OATLD lesions are unlikely to be SCTLD lesions coinfected by *V. coralliilyticus*, a phenomenon found to exacerbate disease progression and mortality by Ushijima et al. (2020). We also assessed the hypothesis that OATLD lesions were the result of amoxicillin treatments on SCTLD affected corals, perhaps selecting for antibiotic resistance and/or more acute lesion progression. Our historical imagery analyses indicated that OATLD was present on SCTLD-treated reefs before such treatments began and was geographically co-occurring with SCTLD. The conclusion driven by our combined results suggests that OATLD is not caused by an amoxicillin-resistant strain of the SCTLD pathogen.

#### White plague/white syndromes

The three white plague types have historically been identified as affecting multiple coral species, while OATLD was only seen on *O. faveolata*. While the upper Keys reefs are largely devoid of other species, some of the lower Keys and inshore reefs where OATLD was observed did contain other white plague-susceptible species, none of which were observed with white plague-like lesions.

White plague has typically been defined as linear, smooth lesions progressing across a colony (Richardson et al., 1998a). This matches the lesion appearance of OATLD. Lesion progression rates were the primary differentiation factor in the descriptions of the three white plague types. Type I was defined as up to 0.3 cm/day, type II up to 2 cm/day, and type III notably exceeding that rate (Dustan, 1977; Richardson et al., 1998a; Richardson et al., 2001). The progression of OATLD lesions, though variable, was overall much faster than type I, but slower than types II and III.

The initiation points of OATLD lesions varied by colony and spread in a variety of directions. White plague type I lesions are similarly documented, but type II lesions are described as almost universally starting at the base of colonies and moving upwards (Richardson et al., 1998a). Descriptions for placement of white plague type III lesions vary, with observations of lesions progressing from the center of colonies (Bythell et al., 2004), as well as from the base (Chaves-Fonnegra et al., 2021).

Prevalence of OATLD displayed a notable seasonal component, peaking during and immediately after peak thermal stress. This coincides with similar observations of white plague (Francini-Filho, 2010; Chaves-Fonnegra et al., 2021).

### Microbiome Assessments

We identified changes in the microbiome associated with OATLD, including the enrichment of specific taxa that may distinguish this disease from SCLTD. Taxa identified as disease specialists were a relatively small proportion of the overall microbiome composition (14%) in *O. faveolata*. Within these specialist taxa, we found taxa that showed similarities and also differences to other known coral diseases.

Among the disease specialists, the OATLD-specific *Aurantivirga* and Micavibrionaceae ASVs had no matches in coral diseases. However, *Cribrihabitans*, *Ferrimonas*, and *Mycoplasma* were each exact sequence matches over the approximately 250-bp V4 region of the 16S rRNA gene to clone library sequences from *O. faveolata* with white plague type II (Sunagawa et al., 2009). Finding exact matches to this previous study is notable given the extensive changes to sequencing technology, analytical tools, and databases in the last fifteen years.

Of these three potential OATLD bioindicators, *Mycoplasma* warrants further investigation. *Mycoplasma* are among the smallest free-living microorganisms and are found in both commensal and pathogenic relationships with diverse animals. *Mycoplasma* has previously been detected in healthy gorgonians (Gray et al., 2011; Holm and Heidelberg, 2016; Quintanilla et al., 2018), the cold-water scleractinian coral *Lophelia pertusa* (now *Desmophyllum pertusum*) (Kellogg et al., 2009; Neulinger et al., 2009), and in several species of jellyfish (Peng et al., 2021; Muffett et al., 2025). Notably, the *Mycoplasma* ASV80 detected here had low similarity (80%) to the *Candidatus* Mycoplasma corallicola from cold-water coral (Neulinger et al., 2009). Applying the 16S rRNA gene similarity thresholds proposed by Yarza et al. (2014), it is likely that the *O. faveolata Mycoplasma* found in the OATLD samples is part of a different bacterial order than *Candidatus* Mycoplasma corallicola.

*Mycoplasma* are naturally resistant to beta-lactam antibiotics, such as amoxicillin, that inhibit cell wall biosynthesis. Since this *Mycoplasma* strain is unique to OATLD compared to SCTLD, and because OATLD is far less responsive to amoxicillin treatments than SCTLD, the role of *Mycoplasma* in OATLD warrants further investigation. Future work should incorporate techniques beyond 16S rRNA studies to evaluate the potential virulence of *Mycoplasma*, such as the development of axenic cultures that would enable experimental transmission studies and the visual confirmation of its presence in OATLD-affected coral tissue via probe-based microscopy.

In addition to OATLD-specific microbial taxa, we detected similarities between OATLD and other coral diseases, including the concurrent SCTLD outbreak in the Florida Keys (Table 1). For example, we detected an altered and more variable microbial community associated with disease lesions, as previously characterized in SCLTD (Meyer et al., 2019; Clark et al., 2021; Huntley et al., 2022), necrotic octocorals (Quintanilla et al., 2018; Keller-Costa et al., 2021), and white plague (MacKnight et al., 2021; Silva-Lima et al., 2021). We also detected an enrichment of the bacterial orders Peptostreptococcales-Tisserales and Rhodobacterales in OATLD lesions that were also more abundant in both SCLTD (Rosales et al., 2020; Rosales et al., 2023) and in white plague type II (Sunagawa et al., 2009). Rhodobacterales were also detected in white plague from *O. annularis* using a microarray of phylogenetic probes, along with members of the polyphyletic Clostridiales which has since been divided into several orders including Peptostreptococcales-Tisserales (Kellogg et al., 2013). Similarity in microbial taxa at the level of bacterial order across multiple coral diseases and host species suggests that these broader groups share functional similarities related to secondary pathogenesis. These opportunistic pathogens may be members of the pathobiome, i.e. normal residents of the coral microbiome that are released from host control in compromised tissues (Sweet and Bulling, 2017). Thus, these broader groups have less utility as bioindicators of specific coral diseases.

### Histology Assessments

Histological examination of OATLD samples showed some similarities and some differences from samples of other described coral diseases (full summary in Supplemental Table 5).

#### SCTLD

Three histological features observed in our OATLD samples are also common in SCTLD-affected corals:

1. Ghosting (necrosis) of endosymbionts in the surface gastrodermis. These could imply potential early stage lesions of the host’s gastrodermal tissue.
2. Necrosis of MFs. We predominantly observed karyorrhexis rather than tissue lysis in the MFs. Presumably, the lesion could expand in association with degrading granular amoebocytes.
3. Depleted and degraded granular amoebocytes (globules) in MFs. These are attributed to lysosomal activity from self-autolysis, potentially leading to rapid tissue necrosis. The depleted (aggregated) granular amoebocytes exhibit a rounded shape that may represent a transitional stage to the degraded state. Similar degraded amoebocytes in the MFs were observed in *Montastraea cavernosa* intentionally infected with SCTLD (Ushijima, unpublished data). It is unclear whether the protrusion of the MFs may contribute to the disease mechanism.

In contrast, most histological characteristics of SCTLD were not observed in the OATLD samples presented here (Table 1). Histopathology of SCTLD-affected corals in Florida identified lytic necrosis of the gastrodermis associated with necrotic, degenerative, deformed endosymbionts (Landsberg et al., 2020; Hawthorn et al., 2024) (Supplemental Figure 4). In contrast, SCTLD-affected corals from the US Virgin Islands exhibited lytic mucus cell hypertrophy in the gastrodermis (Work et al., 2025). In our OATLD samples, neither lytic necrosis nor lytic mucus cell hypertrophy of the gastrodermis was observed. The surface body wall gastrodermis was affected only by foci of necrotic (ghosting) endosymbionts in diseased colonies. Both histology and TEM identified swollen or larger symbionts in OATLD-affected corals, which is a contrast to SCTLD studies in which endosymbionts were atrophied, vacuolated, misshapen, and necrotic (Landsberg et al., 2020; Hawthorn et al., 2024). In all of these respects, OATLD does not fit the current case definition of SCTLD.

#### White plague/white syndromes

Early histopathology reports on white plague types I and II revealed necrosis at the tissue loss margins, tissue degradation, and degenerating zooxanthellae in the surface body wall gastrodermis (Peters, 1984; Richardson et al., 1998b; Bythell et al., 2002). White plague type III-affected corals were not examined microscopically.

Samples from an outbreak of white syndrome across multiple species in Flower Garden Banks National Marine Sanctuary in October 2022 and March 2023 were histologically assessed by Rossin (2024). Assessments focused primarily on microscopic lesions in the surface body wall of the gastrodermis; the condition of deeper polyp tissues was not reported. Primary findings were necrosis (“ghosting”) and degradation in the endosymbionts, vacuolization, exocytosis, and separation of the gastrodermis from the mesoglea. The study concluded that diseased samples showed intensive necrosis and consistent exocytosis and degraded symbionts, but that the lack of vacuolization in the symbionts, a defining characteristic of SCTLD, indicated that the outbreak was “likely more similar to white plague.” Though the mesenteries and mesenterial filaments were not examined by Rossin (2024), many of the histological conditions observed from Flower Garden Banks are similar to those seen in our OATLD samples.

Samples from *O. faveolata* affected with an unknown white-syndrome (along with healthy controls) collected in 2004 to 2006 from five sites in the western tropical Atlantic underwent a preliminary histopathological examination (Peters, unpublished data). Some of these samples exhibited lytic necrosis of the gastrodermis and similar other changes seen later in SCTLD-affected corals. A set of samples exhibiting an unknown white disease were also collected from Dry Tortugas on Florida’s Coral Reef in 2016 (5 years prior to the arrival of SCTLD). These samples were also taken from both diseased and healthy colonies, but of the congener *O. franksi* (Peters, unpublished data: Table 1, Supplemental Table 5). Among these, most healthy samples had mild gastrodermal changes but normal endosymbionts; however, one apparently healthy sample had suspect magenta coccobacilli in the gastrodermis. The diseased samples exhibited ghosting endosymbionts, gastrodermal mucocyte hypertrophy, and necrosis of gastrodermal cells in the mesenteries attached to the cnidoglandular bands, similar to our OATLD-affected colonies. However, lytic necrosis of the gastrodermis was not observed in the *O. franksi* samples.

### TEM Assessments

OATLD-affected coral tissues typically had fewer, but larger, symbionts than non-diseased tissues. This may reflect a trade-off for tissues needing to regulate endosymbiont size and density. Greater endosymbiont densities have been negatively correlated to symbiont size, and nutrient availability may play a role in this trade-off (Jones and Yellowlees, 1997). It is possible that OATLD may create a nutrient-limiting environment within affected tissues.

Symbionts from OATLD-affected coral tissues contained more and larger lipid droplets than adjacent non-diseased samples. A similar pattern was true for starch granules, with symbionts from D samples having a greater number and a larger area than H samples (with U samples having intermediate values). Symbionts from diseased tissues may be storing starch and lipid reserves in preparation of being expelled; this could be a strategy allowing for survival in the water column where efficient photosynthesis may be hindered. The increase in lipid and starch stores could also indicate that OATLD-affected coral tissues and/or symbionts are not utilizing these reserves as readily as non-affected tissues. Increased starches in Symbiodiniaceae have also been attributed to nitrogen starvation (Ishii et al., 2025) as well as host starvation combined with light depletion (Muller-Parker et al., 1996).

Symbionts from healthy corals had a higher prevalence of electron dense bodies than samples from OATLD-affected corals. Electron dense bodies appear dark in micrographs and could encompass organelles such as lysosomes, viral components, or nucleic acids. It is unclear how this relates to health states, particularly as only a few symbionts contained these electron dense bodies. Further investigation is needed to determine the composition and role of electron dense bodies, particularly regarding coral disease signs.

Degradation of the symbionts or their organelles was not a significant parameter between health states. Though cellular degradation is a general indicator for necrosis or apoptosis, it may not be observed in these samples because of the rapid rates of tissue loss associated with OATLD. The numerous other parameters assessed (including cavities, accumulation bodies, membrane separation, pyrenoids, and VLPs) did not vary among health states. Though further investigation into OATLD samples could yield additional information, it is possible that the causative agent(s) do not impact these ultrastructure features.

#### SCTLD

Our direct comparisons between OATLD assessments and a previous SCTLD dataset identify some differences in cellular pathology (Table 1). For some parameters, these differences were not observed when only *O. faveolata* colonies were compared, but were observed in comparisons of the multi-species SCTLD dataset.

Specifically, both starch and lipid stores were increased in OATLD-diseased samples compared to apparently healthy tissues. In SCTLD-affected corals, starch and lipid stores decreased. This could indicate physiological differences that occur during these two potentially distinct diseases, which could include changes to symbiont health, photosynthetic activity or efficiency, or carbon exchange between the coral and the symbiont.

Other metrics, such as chloroplast gigantism, the presence of cavities, pyrenoid presence, and the presence of a starch sheath around the pyrenoid were not associated with diseased OATLD samples, but are previously correlated with SCTLD.

Viral-like particles are prevalent in SCTLD samples and have been correlated to the diseased health state (Work et al., 2021), but it is unclear whether they are involved in SCTLD pathogenesis. VLPs were also prevalent in all OATLD health states, and were not an indicator of disease. Contrasting SCTLD findings, the prevalence of electron dense bodies was not common in the OATLD samples regardless of health state. In SCTLD corals, prevalence of electron dense bodies was higher in diseased corals, while in OATLD corals, it was higher in healthy corals (6/10 total samples with this parameter). Additional studies assessing the role of VLPs and electron dense bodies in pathogenesis are needed for both OATLD and SCTLD, as this cannot be determined through bioimaging alone.

#### White plague/white syndromes

Though the use of TEM has become increasingly common in coral disease studies, there are very few descriptions of TEM micrographs for white syndromes. Tissue and symbiont degeneration were observed with TEM by Richardson et al. (1998a) in corals affected by white plague II, but no pathogens were observed. Denner et al. (2003) conducted TEM examinations of white plague II corals to describe the subsequently identified pathogen, but did not assess damage to coral tissue or symbionts. Soffer et al. (2014) used TEM to document degraded tissue, and numerous viral-like particles in white plague II tissues, though they did not link the presence of the viruses to disease state. With these limited studies and lack of quantifiable tissue and endosymbiont metrics, it is difficult to assess how TEM descriptions of other white diseases compare to our OATLD observations.

### Synthesis and Recommendations

The differences in gross appearance, lesion behavior, microbiome composition, histopathological parameters of both coral tissues and endosymbionts, and physiological metrics of endosymbionts suggest that OATLD is likely a different disease from SCTLD (Table 1). Though there are distinct differences between the two, the similarities that do exist in microbiome components as well as physiological responses of the coral and symbionts affected by both types of lesions provide future opportunities to explore what characteristics may be present across multiple diseases as compared to those associated with any specific disease.

The comparison of OATLD to white plague or other white syndromes is less clear (Table 1). The most notable differences of OATLD from definitions of white plague are 1) the occurrence on only *O. faveolata* and not on any other known susceptible species and 2) the movement of lesions in any direction across affected colonies. Our laboratory analyses can neither confirm nor refute a distinction between the diseases. This is largely because there is a dearth of histology or TEM on white plague against which to compare our OATLD findings. Our histological comparisons to presumed white plague in *O. franksi* provide a starting point, but we suggest that the field would be significantly enhanced by quantitative and qualitative histological and TEM analyses of each classic white plague type on *O. faveolata* and the subsequent development of case definitions. In the absence of distinctions aside from those seen in situ, it is possible that our description here of OATLD is in fact a definition of a species-specific white plague.

Our analyses highlight some areas for further study of coral and endosymbiont physiology, particularly within disease lesions. The role of VLPs and electron dense bodies within corals and endosymbionts remains largely unknown, and our analyses here add complexity to this avenue of research. One interesting, and potentially novel, physiological response to disease observed here was the changes to the mesenterial filaments. We suggest that further study on the ultrastructural changes of those, along with searching for microorganisms within those areas, may provide additional insights. Within the microbiome, *Mycoplasma* emerged as a taxon of interest that was not previously associated with coral disease.

We also recommend further work on OATLD. Initial experiments should focus on whether the disease is transmissible, both among *O. faveolata* colonies as well as to other species. These experiments would also allow opportunities to assess whether mesenterial filaments are exhibiting unusual behavior (whether they protrude normally at nighttime), as well as to conduct time-series sampling to follow disease progression. We also highly recommend the development of treatment strategies to halt OATLD lesions. As of autumn 2025, OATLD continues to affect many of the largest surviving corals in Florida. Recent successes in developing in situ treatment options for diseases (Randall et al., 2018; Neely et al., 2021b; Eaton et al., 2022; Pitts et al., 2025) suggests that mortality from OATLD may be preventable should intervention options be trialed and permitted.

The collaborative nature of this project provided valuable cross-disciplinary understanding and quantification of disease metrics. We suggest that the integration of field observations and experiments, microbiome analyses, and microscopic examination at multiple scales of both coral hosts and endosymbionts can provide a more complete picture of disease etiology. Each component represents a powerful tool and can provide quantitative metrics for comparison, and together highlight trends and patterns across the coral holobiont. We suggest that such collaborative analyses are likely to create more cohesive understandings of coral stressors.

## Acknowledgements

Field monitoring occurred under an amendment to Sanctuary permit FKNMS-2020-077. Sampling was conducted under FKNMS-2023-141-A1. We are grateful to Kathryn Toth for assistance in fieldwork and historical imagery analyses. We thank the state of Florida for funds under the Florida Fish and Wildlife Conservation Commission’s (FWC) Fish and Wildlife Research Institute’s (FWRI) Fish and Wildlife Health Program, and are grateful to Noretta Perry, Mathew Myers, Yvonne Waters, and Patrick Wilson at FWC/FWRI for preparing the histology slides. Imaging data was collected using the Richard M. Dillaman Biological Imaging Facility at the University of North Carolina Wilmington, and we thank Dr. Alison Taylor and Elizabeth Elliott for technical assistance with TEM imaging.

## Supplemental Figures

**Supplemental Figure 1.**
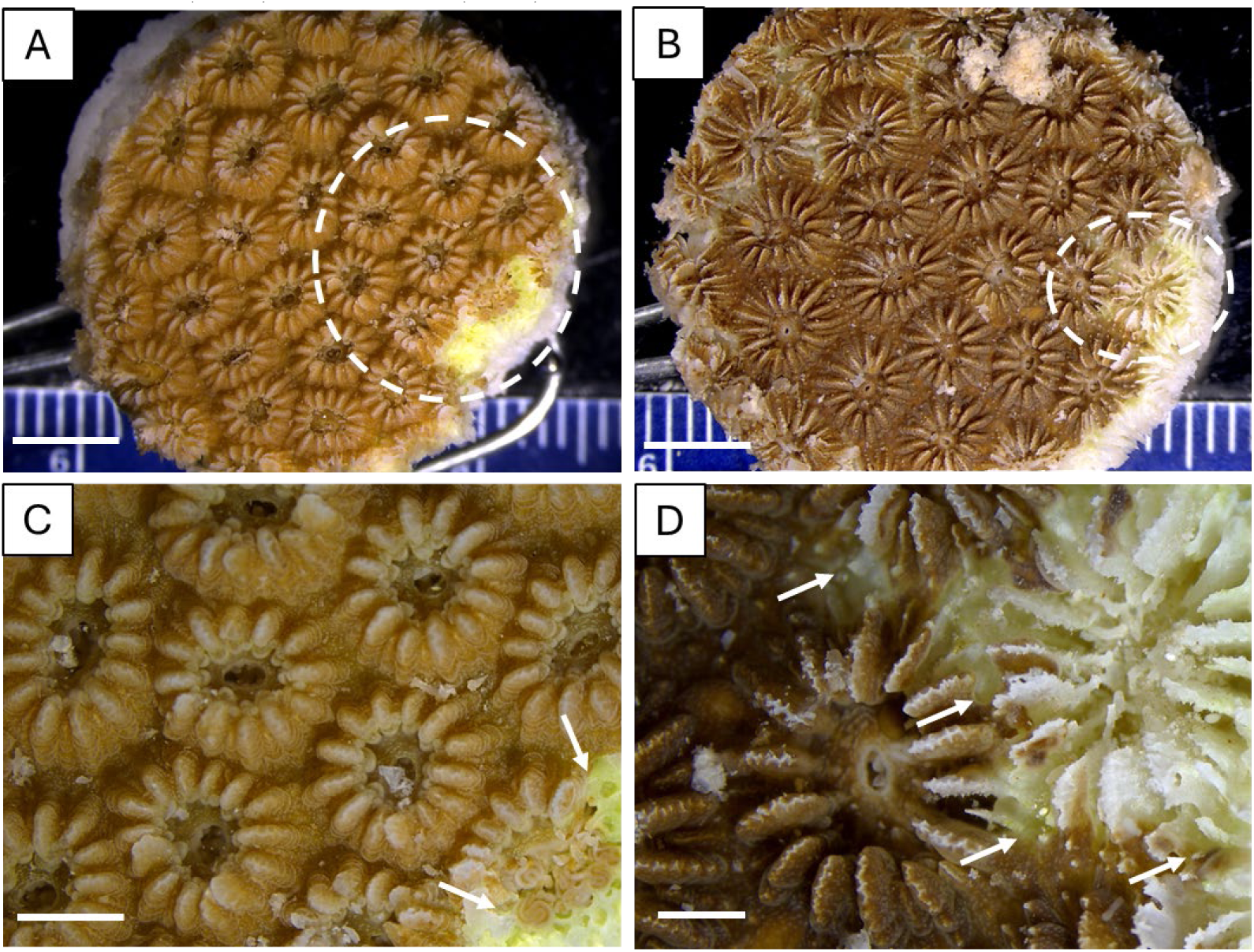
Macrophotographs of two fixed diseased *Orbicella faveolata* tissue samples. Four of the 15 D samples exhibited this type of gross lesion. **(A,B)** Dotted circles highlight areas of possible tissue loss at the edge of the samples. **(C, D)** High magnification views of the circled areas, showing the border of tissue loss and healthy tissue (arrows). Scale bars: 5 mm (A-B) and 2 mm (C-D).

**Supplemental Figure 2.**
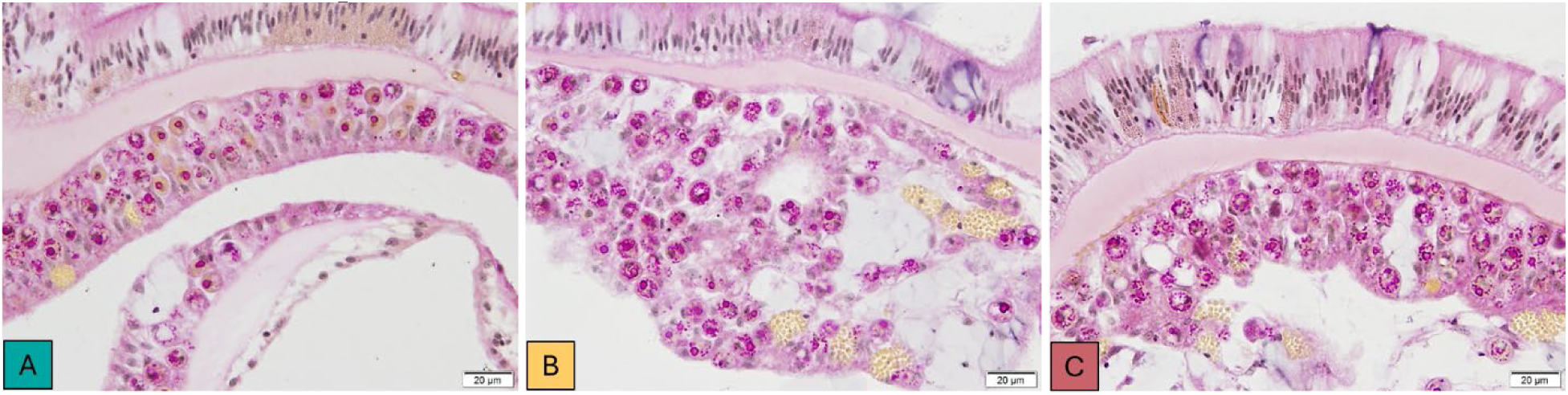
Histological sections of the gastrodermis surface body wall of *Orbicella faveolata* showing endosymbiont starch granules with strong periodic acid-Schiff (PAS)-positive reaction (PAS-MY). (**A**) Healthy sample. (**B**) Disease unaffected sample. (**C**) Disease affected sample.

**Supplemental Figure 3.**
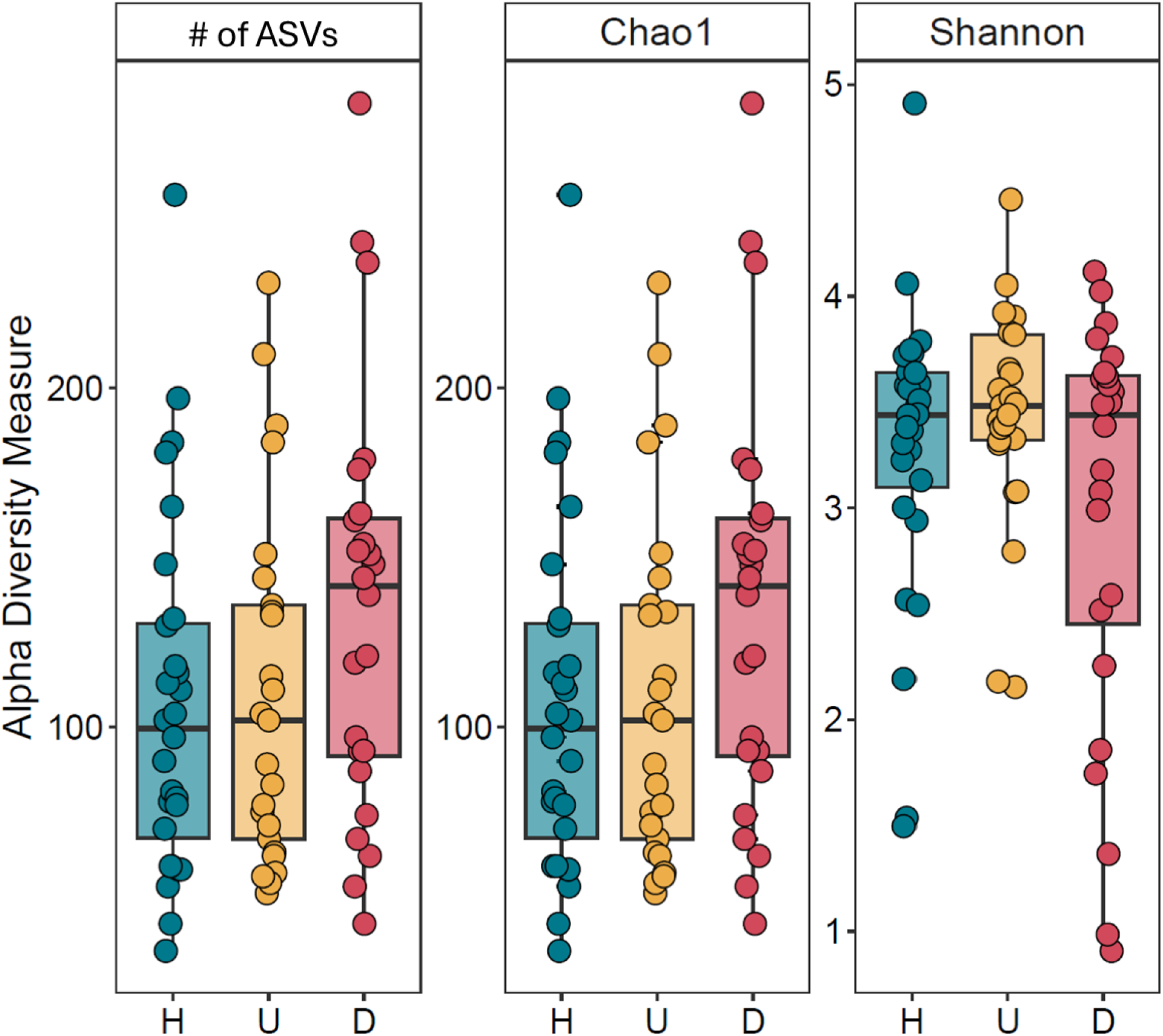
Alpha diversity metrics in microbiomes from *Orbicella faveolata* coral colonies, including the observed number (richness) of Amplicon Sequence Variants (ASVs), Chao1 diversity, and Shannon diversity. Microbiome samples were sourced from healthy colonies (H), unaffected areas of diseased corals (U), and disease lesions (D).

**Supplemental Figure 4.**
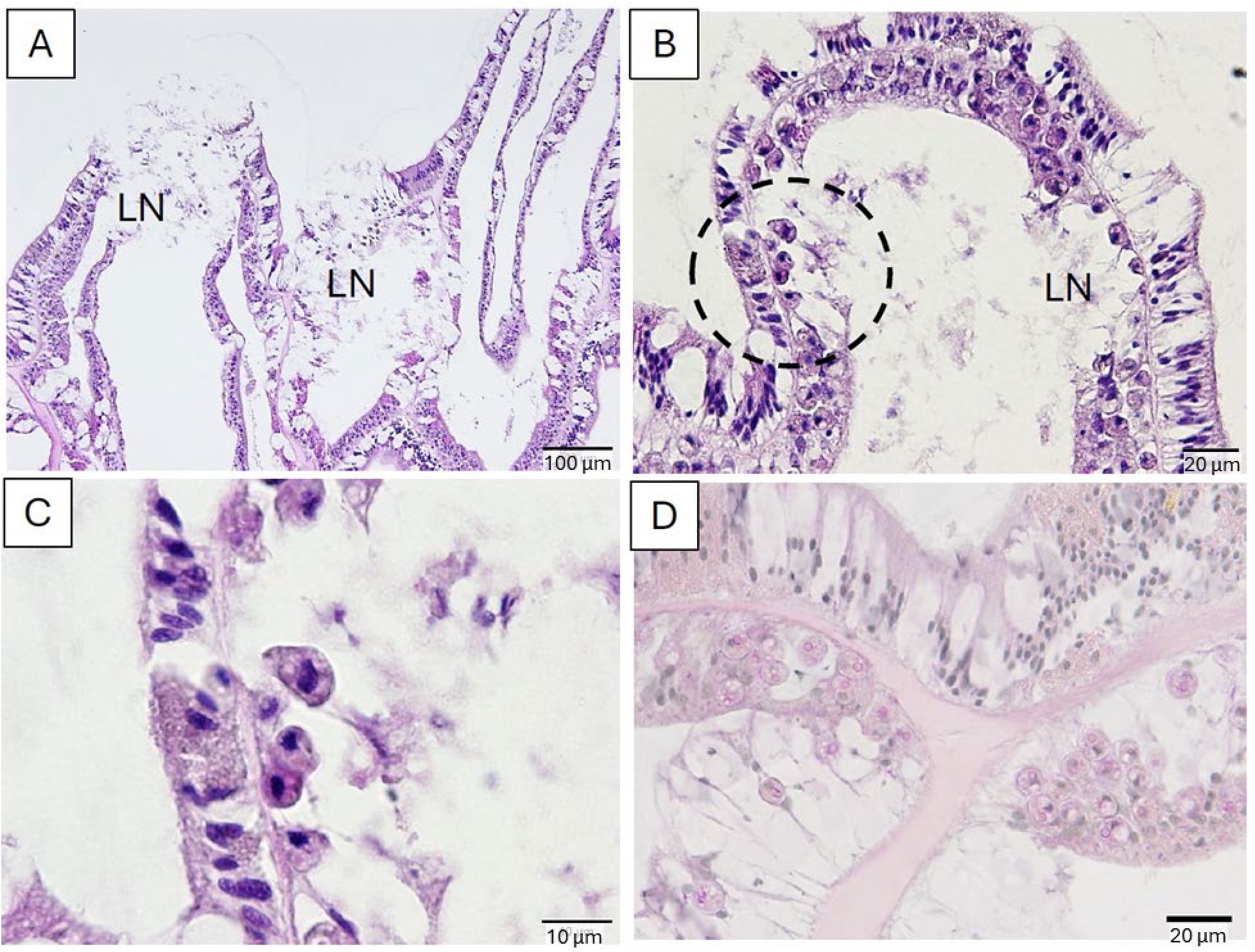
Histological sections of SCTLD-affected *Orbicella faveolata* for comparative pathology. Samples were collected in 2018 in the Florida Keys (A-C stained with H&E; D stained with PAS-MY). (**A**) Lytic necrosis (LN) is shown in the gastrodermis (both basal and surface body wall) and beyond the epidermis. The granular amoebocytes in the mesenterial filaments are depleted (granular amoebocytes from healthy samples were abundant; see Figure 8F). (**B**) Another example of LN of the surface body wall gastrodermis. (**C**) High magnification of dashed circled area in (B) showing deformed, necrotic endosymbionts. (**D**) Endosymbionts in the surface body wall gastrodermis exhibited PAS lightly positive reaction.

